# A conserved chronobiological complex times *C. elegans* development

**DOI:** 10.1101/2024.05.09.593322

**Authors:** Rebecca K. Spangler, Kathrin Braun, Guinevere E. Ashley, Marit van der Does, Daniel Wruck, Andrea Ramos Coronado, James Matthew Ragle, Vytautas Iesmantavicius, Lucas J. Morales Moya, Keya Jonnalagadda, Carrie L. Partch, Helge Großhans, Jordan D. Ward

## Abstract

The mammalian PAS-domain protein PERIOD (PER) and its *C. elegans* orthologue LIN-42 have been proposed to constitute an evolutionary link between two distinct, circadian and developmental, timing systems. However, while the function of PER in animal circadian rhythms is well understood molecularly and mechanistically, this is not true for LIN-42’s function in timing rhythmic development. Here, using targeted deletions, we find that the LIN-42 PAS domains are dispensable for the protein’s function in timing molts. Instead, we observe arrhythmic molts upon deletion of a distinct sequence element, conserved with PER. We show that this element, designated CK1δ-binding domain (CK1BD), mediates stable binding to KIN-20, the *C. elegans* CK1δ/ε orthologue. We demonstrate that CK1δ phosphorylates LIN-42 and define two conserved helical motifs in the CK1BD, CK1BD-A and CK1BD-B, that have distinct roles in controlling CK1δ-binding and kinase activity *in vitro*. KIN-20 and the LIN-42 CK1BD are required for proper molting timing *in vivo*, and loss of LIN-42 binding changes KIN-20 subcellular localization. The interactions mirror the central role of a stable circadian PER–CK1 complex in setting a robust ∼24-hour period. Hence, our results establish LIN-42/PER – KIN-20/CK1δ/ε as a functionally conserved signaling module of two distinct chronobiological systems.

## Introduction

Chronobiology is the study of biological rhythms. Circadian rhythms which enable organisms to anticipate daily cycles of light, temperature, and other environmental variables are one of the best studied examples. ^1^ PERIOD (PER) proteins are central to this timekeeping mechanism. Their cellular abundance, stability, and nuclear localization changes rhythmically over a ∼24-hour cycle. In mammals, the paralogs PERIOD1 and PERIOD2 (PER1 and PER2) associate with Casein kinase 1 δ/ε (CK1δ/ε) as well as the CRYPTOCHROME proteins, CRY1 and CRY2, to form nuclear transcriptional complexes which repress the transcriptional activators circadian locomotor output cycles protein kaput (CLOCK) and brain and muscle ARNT-like 1 (BMAL1)(Fig. S1A).^2–5^ Since the targets of the CLOCK-BMAL1 heterodimer include PER and CRY themselves, PER and CRY thus eventually repress their own production, permitting subsequent re-expression of CLOCK-BMAL1 to start a new cycle.^2,6^

To maintain the 24-hour period of circadian rhythms, a delay between the activation and the repression of CLOCK-BMAL1 transcription is required. Central to this delay is the post-translational modification of PER by one of the two closely related CK1δ and CK1ε kinases. For convenience and because of the redundancy in their function, we will henceforth refer to the two isoforms generically as CK1. Anchored through a CK1-Binding Domain (CK1BD)^7^, CK1 is associated with PER for its entire existence in the cell, even translocating from the cytoplasm to the nucleus together (Fig. S1A)^4,8^, allowing PER to deliver CK1 to other targets at clock-controlled promoters.^8,9^ CK1-mediated phosphorylation of a PER degron licenses PER ubiquitylation and subsequent degradation, thereby limiting PER abundance and its period of activity (Fig. S1B).^10–12^ In addition to the degron, CK1 phosphorylates multiple serines in another PER sequence element known as FASP (“Familial Advanced Sleep Phase”)(Fig. S1B).^13,14^ Phosphorylation of the FASP site turns PER into an inhibitor of CK1 kinase activity.^15,16^ In humans, a single residue mutation in this region that blocks all FASP phosphorylation results in a short circadian period that manifests as the eponymous Familial Advanced Sleep Phase, where affected individuals wake up very early in the morning.^17,18^

In contrast to circadian timing, the chronobiology of development is mechanistically less well understood. In *C. elegans*, a single PER orthologue, LIN-42, appears to diverge substantially in structure and function from the mammalian and fly PER proteins. At 598 amino acids for its longest isoform, LIN-42 is substantially shorter than mouse PER2 at 1225 amino acids.^19^ LIN-42 lacks a CRY-binding domain, consistent with a lack of a CRY orthologue in *C. elegans*, and in its tandem PER-ARNT-SIM (PAS) domains, only PAS-B is well conserved, adopts a canonical fold, and mediates dimerization in a mode identical to mammalian PER.^20,21^ Finally, two stretches of sequence, previously termed SYQ and LT according to their first amino acids^22^, bear sequence homology to the two PER CK1BD subdomains, CK1BD-A and CK1BD-B, but their function has remained unexplored. *lin-42* further differs from canonical PERs in its expression dynamics and functions: rather than exhibiting a ∼24-hour, temperature-invariant period, *lin-42* expression cycles exhibit a temperature-dependent length which can be as short as ∼7 hours at 25°C.^19,23^ Indeed, *lin-42* was identified as a developmental gene whose mutation causes heterochronic phenotypes, i.e., defects in temporal cell fate specification where cells adopt adult cell fates precociously, in larvae.^24,25^ Hence, it was proposed that *lin-42/PER* constitute an evolutionary link between two distinct, circadian and developmental, timing systems.^19^

The extent of functional similarity between these timing systems generally, and LIN-42 and PER function specifically, has remained uncertain, despite the realization that additional orthologues of mammalian clock genes exist and cause heterochronic phenotypes when mutated.^19,21,23,26–28^ Conceptually, whereas mutations of circadian clock genes such as *PER* change the tempo and/or robustness of circadian rhythms, heterochronic mutations are defined by their ability to alter the sequence of developmental events, such that certain events are skipped.^29,30^ Accordingly, and despite its rhythmic expression, the heterochronic function of LIN-42 does not appear to involve recurring activity but rather the stage-specific repression of the *let-7* miRNA prior to the third larval stage.^31–33^

LIN-42 is also required for the rhythmic occurrence of molts. Wild-type animals molt, i.e., regenerate a new collagenous apical extracellular matrix (cuticle), at the end of each of the four larval stages.^34^ Under constant environmental conditions, individually grown animals enter and exit molts with great temporal uniformity, revealing robust temporal control. *lin-42(ok2385)* mutant animals were shown to cause a slow-down of development as well as an arrhythmic molting phenotype where this uniformity is lost such that individual animals molt at different times.^35,36^ It is unknown how LIN-42 contributes to rhythmic molting mechanistically. In addition to *CRY*, the *C. elegans* genome lacks obvious orthologues of *BMAL1* and *CLOCK*, arguing for a mechanism that differs from that of the circadian clock at least in the identity of several core components.^27^ In a yeast two-hybrid assay, LIN-42 was shown to be capable of binding to numerous transcription factors.^37^ Although the functional relevance of such binding for molting has remained unexplored, LIN-42 binding to the REV-ERB orthologue NHR-85 appears important for robust periodic transcription of the heterochronic *lin-4* miRNA. Finally, loss of KIN-20, the *C. elegans* orthologue of CK1δ/ε, replicates some heterochronic phenotypes of *lin-42* mutation and slows development^26^, but its heterochronic functions were argued not to involve LIN-42^38^, and it is unknown whether KIN-20 is required for rhythmic molting.

Here, we set out to further characterize the molecular and developmental functions of LIN-42. Using targeted mutations, we find that its PAS domains are largely dispensable both for heterochronic pathway activity and rhythmic molting. By contrast, the SYQ/LT regions are specifically required for rhythmic molting. We demonstrate that these domains function as a CK1BD, mediating stable KIN-20/CK1 binding to LIN-42 *in vivo* and *in vitro*. CK1 phosphorylates LIN-42 *in vitro* and this activity is distinctly modulated through the two CK1BD subdomains: CK1BD-B is particularly important for stable CK1 binding, whereas the CK1BD-A element inhibits CK1 enzymatic activity. Moreover, loss of LIN-42 binding impairs nuclear localization of KIN-20. Our results identify LIN-42/PER–KIN-20/CK1 as a conserved chronobiological complex utilized by two distinct biological oscillators. Although CK1 activity in the circadian clock has previously been viewed mostly through the lens of its effects on PER, our findings align well with the growing notion that PER-mediated regulation of CK1 may be an additional important mechanism to support robust circadian rhythms.

## Results

### The LIN-42 CK1BD mediates a subset of developmental functions of LIN-42

Previous analysis of the two partial deletion alleles of *lin-42*, *n1089* and *ok2385* (Fig. 1A), have suggested the possibility that different developmental functions could be genetically separable. Although both deletions caused precocious heterochronic phenotypes, only *ok2385* showed evidence of slow and arrhythmic molting.^22,35,39^ To confirm these findings, we quantified molt, intermolt, and larval stage duration for mutant and wild-type animals at high temporal resolution, using a luciferase assay that monitors entry and exit from the lethargus state associated with molting on animals grown in isolation.^36,40^ Indeed, we found that *lin-42(ok2385)* mutant larvae developed more slowly than wild-type animals and became increasingly arrhythmic (Fig. 1B, S2A-C). Moreover, and as observed previously^36^, most mutant animals underwent only three molts (33/39 animals) within the duration of this assay, suggesting either a precocious exit from the molting cycle, or a very delayed or abnormal fourth molt (Fig. 1B, S2D). By contrast, essentially all *lin-42(n1089)* animals completed a normal number of molts (42/42 animals) and maintained robust synchrony, although they developed significantly slower than wild-type animals (Fig. 1B). At the same time, both alleles caused robust heterochronic phenotypes, illustrated by precocious formation of alae (Fig. 1C) in L3 to early L4 larvae. Alae are a cuticular structure normally secreted by the terminally differentiated epidermal seam cells at the L4-adult transition.

**Figure 1.**
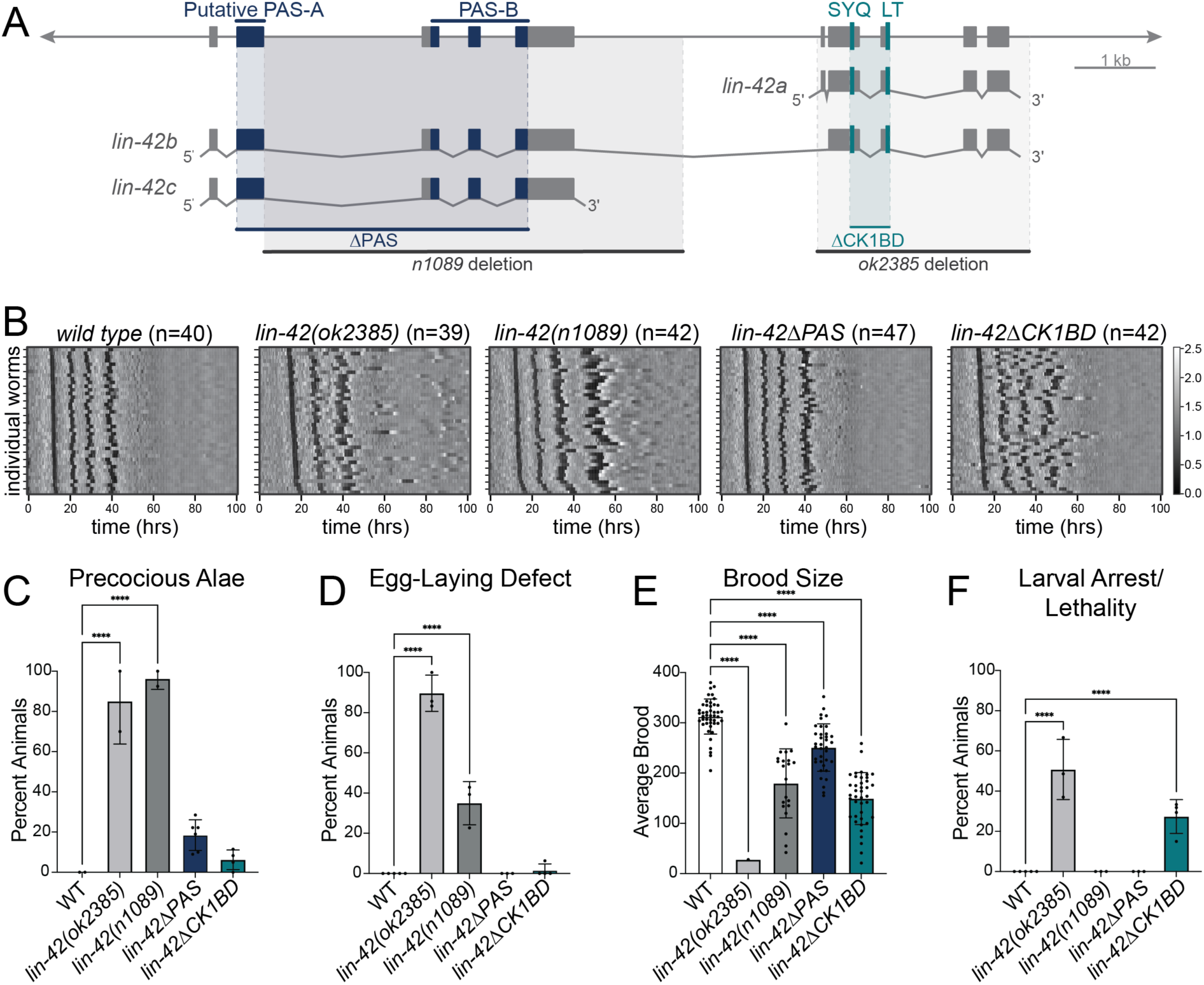
*lin-42ΔPAS* mutations cause mild heterochronic phenotypes while *lin-42ΔCK1BD* mutations cause asynchronous molting. **(A)** Schematic depiction of the *lin-42* genomic locus. The gene structure of the *lin-42 a, b*, and *c* isoforms are shown. The location of sequences encoding putative PAS-A, PAS-B, SYQ, and LT sequence motifs are indicated. Published *lin-42* deletion alleles *n1089* and *ok2385* are indicated in dark and light grey, respectively, newly generated targeted *ΔPAS* and *ΔCK1BD* deletions in navy and teal, respectively. **(B)** Heatmaps showing trend-corrected luminescence traces from the indicated genotype. Each horizontal line represents one animal. Traces are sorted by entry into the first molt. Darker color indicates low luminescence signal and corresponds to the molts. **(C)** Bar plot quantifying the percentage of animals of the indicated genotype with precocious complete or partial alae at the L3-L4 molt. **(D)** Bar plot quantifying the percentage of animals of the indicated genotype that exhibited egg-laying defects as determined by the presence of hatched larvae in the animal. **(E)** Bar plot depicting the average number of live progeny from hermaphrodites of the indicated genotype. **(F)** Bar plot quantifying the percentage of animals of the indicated genotype that arrested or died as larvae. (C-F) Statistical significance was determined using an ordinary one-way ANOVA. p<0.05 was considered statistically significant. **** indicates < 0.0001. Error bars in C-F represent standard deviation.

The complex nature of the alleles, each with at least one breakpoint in a noncoding region, precludes a straightforward attribution of the phenotypes to specific features of the LIN-42 protein. Therefore, we decided to generate precise deletions of the two conserved regions, PAS and SYQ/LT (Fig. 1A). Henceforth, and based on the functional characterization which we describe below, we will refer to SYQ/LT as CK1BD for CK1-Binding Domain. To our surprise, *lin-42(wrd67[ΔPAS])* mutant animals exhibited essentially wild-type molting patterns in the luciferase assay, resembling neither *lin-42(n1089)* nor *lin-42(ok2385)* mutant animals. By contrast, *lin-42(wrd63[ΔCK1BD])* mutant animals exhibited increasing loss of molting synchrony over time (Fig. 1B). However, unlike *lin-42(ok2385)*, *lin-42(ΔCK1BD)* mutant animals executed four detectable molts during the assay (Fig. 1B, S2D).

To get a better understanding of the developmental relevance of each domain, we examined additional phenotypes (Fig. 1D-F; Table S1). We found that all four alleles affected brood sizes, albeit to varying degrees, in the order *ok2385>>ΔCK1BD≈n1089>ΔPAS* of decreasing severity (Fig. 1E, Table S1). Only *ok2385* and *n1089* caused highly penetrant egg-laying (Egl) and precocious alae defects^22,35,39^, whereas ΔPAS and ΔCK1BD mutants had completely wild-type egg laying and only modest precocious alae (Fig. 1C and D, Table S1). Finally, both *ok2385* and, to a lesser extent, ΔCK1BD caused larval arrest and lethality, a phenotype interestingly not observed in the luciferase assay, suggesting an environmental dependency (Fig. 1F, Table S1). Taken together, these results indicate surprisingly mild phenotypes upon loss of the LIN-42 PAS region and highlight an important function of the CK1BD for rhythmic molting.

### LIN-42 and KIN-20 interact *in vivo*

Since the genetic results suggested separable functions of LIN-42, we hypothesized that these could depend on specific interaction partners. To identify interaction partners we performed anti-FLAG immunoprecipitations on endogenously tagged *lin-42(xe321[3xFLAG::lin-42b+c])* mixed-stage animal lysates followed by mass-spectrometry. As a control for non-specific binding, we also performed an anti-FLAG immunoprecipitation in an unrelated strain (*sart-3::GFP::3xFLAG*).^41^ We observed a single highly enriched interaction partner for LIN-42: KIN-20 – the orthologue of mammalian CK1δ/ε (Fig. 2A).

**Figure 2.**
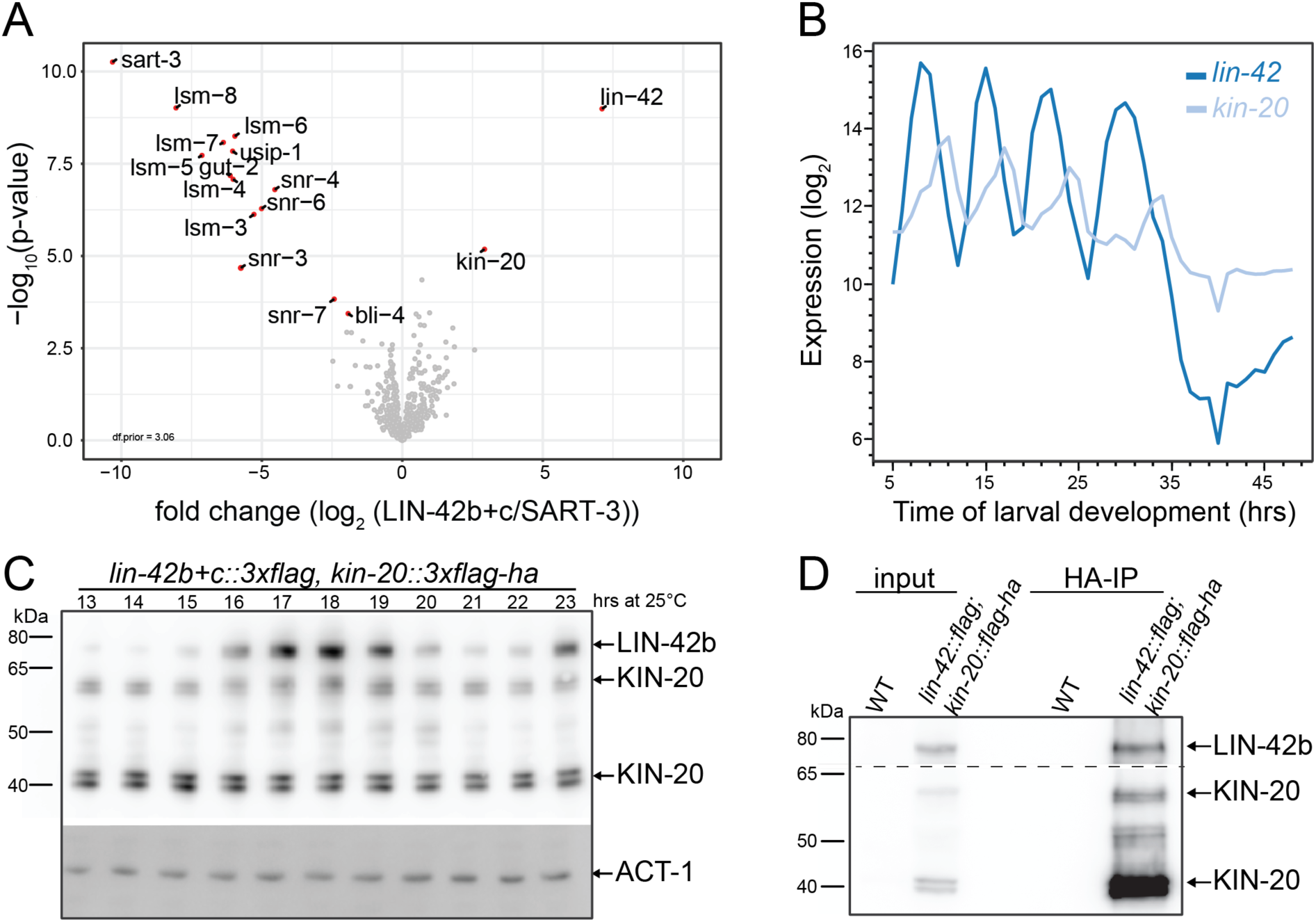
LIN-42 and KIN-20 interact *in vivo*. **(A)** Volcano blot comparing protein enrichments in 3xFLAG::LIN-42b+c and 3xFLAG::SART-3 (control) immunoprecipitations, determined by mass spectrometry. **(B)** mRNA expression profile for *lin-42* (dark blue) and *kin-20* (light blue) mRNAs throughout larval development. Data Source: Meeuse et al., 2020. **(C)** Western blot with extracts from *3xflag::lin-42b+c; 3xflag-ha::kin-20* (HW3479) larvae collected hourly for 11 hours, starting at 13 h after plating synchronized L1 stage animals on food at 25°C. Top part probed with anti-FLAG-HRP (1:1,000), lower part with anti-actin-1 (1:7,500). Arrows indicate bands for LIN-42b, KIN-20 and ACT-1 (see Fig. S5A). **(D)** Anti-HA pulldown from *3xflag::lin-42b+c; 3xflag-ha::kin-20* animals. Blot probed with anti-FLAG (1:1,000).

The mRNA levels of both *lin-42* and *kin-20* oscillate.^40,42^ Further analysis of the published sequencing data revealed that the peak phases of the two proteins differ, but that there are periods where both transcripts accumulate (Fig. 2B). This is also true for the proteins: we observed by Western blotting that the levels of endogenously tagged KIN-20 protein were largely invariant over time, whereas those of endogenously tagged LIN-42b protein levels changed rhythmically (Fig. 2C). Finally, we confirmed a physical interaction with a reciprocal anti-HA pulldown from *lin-42(xe321[3xFLAG::lin-42b+c]); kin-20(xe328[3xFLAG-HA::kin-20])* animal lysates, again using endogenously tagged proteins (Fig. 2D). Taken together, we conclude that LIN-42 and KIN-20 form a complex *in vivo*.

### The LIN-42 SYQ-LT regions form a functional CK1BD capable of binding mammalian CK1

Mammalian CK1 binds PER through its kinase domain. This region is highly conserved, with CK1δ and KIN-20 exhibiting 79% and 100% sequence identity within the kinase domain and active sites, respectively (Fig. S3).^43,44^ By contrast, the CK1BD is less well conserved in LIN-42 with only ∼30% sequence identity and a different spacing of the relative LIN-42 CK1BD-A and -B sites (Fig. 3A). Hence, it was unclear whether this region could function as a CK1BD in LIN-42.

**Figure 3.**
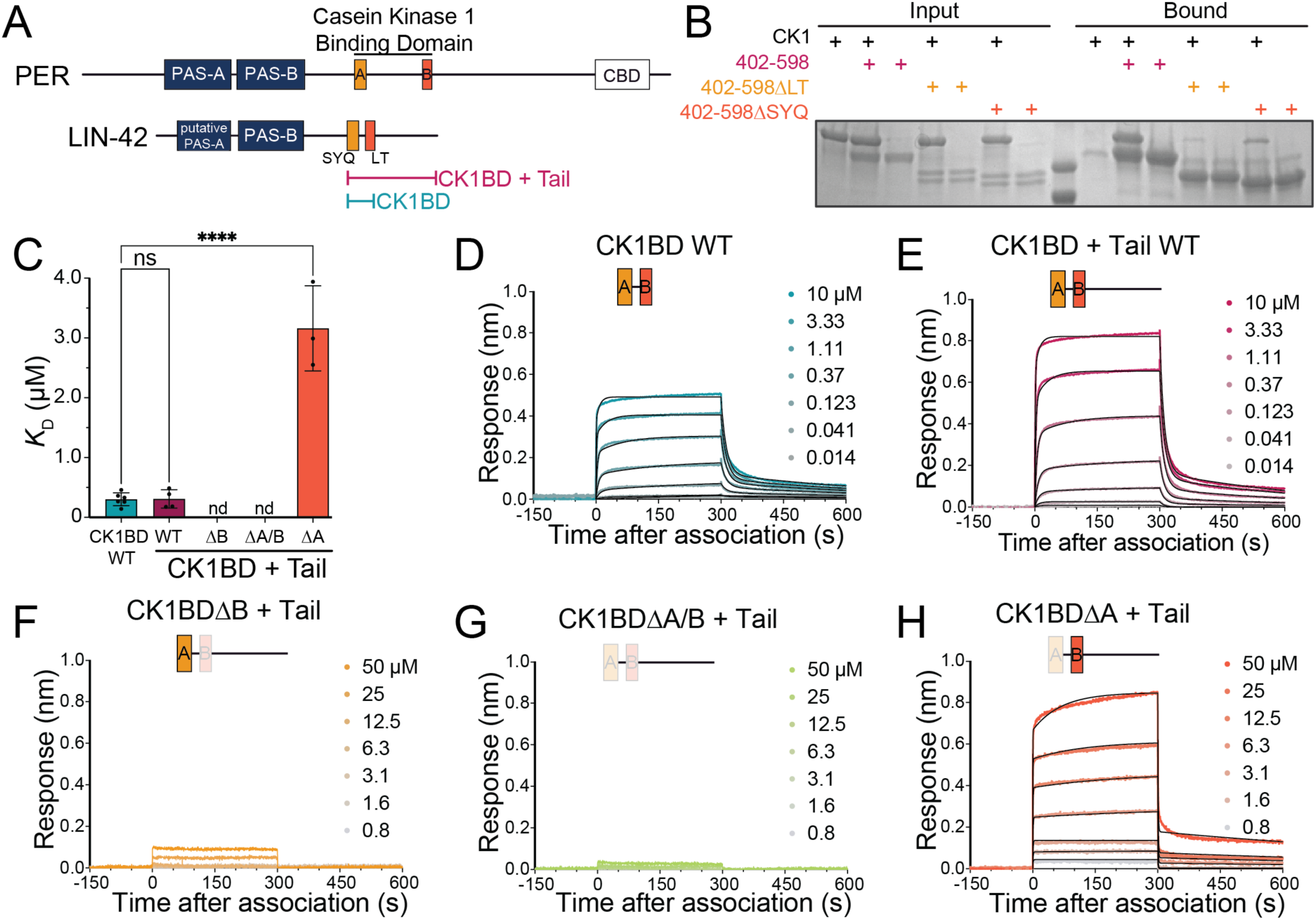
The LIN-42 SYQ and LT domains constitute a functional CK1-binding domain. **(A)** Schematic representing PER2 and LIN-42 protein domains. Protein constructs used in this study, CK1BD + Tail and CK1BD, are indicated i. CK1BD, Casein Kinase 1-Binding Domain; CBD, CRY-binding domain. **(B)** Representative pulldown assay of human CK1 and biotinylated LIN-42 CK1BD + Tail proteins using the indicated protein variants. **(C)** Values for *K*_D_ from kinetic analysis of BLI data (D-H) based on a 2:1 heterogeneous ligand binding model and global analysis (Octet). Mean ± SEM, ordinary one-way ANOVA. p<0.05 was considered statistically significant. **** indicates p< 0.0001. **(D-H)** Bio-layer interferometry (BLI) data for indicated LIN-42 protein binding to immobilized, biotinylated CK1. Inset values represent the concentrations of LIN-42 for individual binding reactions. Model fit to association and dissociation over time is represented by black lines. Data shown from one representative experiment of n ≥ 3 assays.

We produced recombinant proteins to test this possibility. We were unable to express soluble KIN-20 from a bacterial or insect system, so we used recombinant human CK1δ kinase domain (hereafter referred to as CK1), similar to previous studies of KIN-20.^45^ *In vitro* pulldown assays with biotinylated LIN-42 protein containing the SYQ/CK1BD-A and LT/CK1BD-B domains as well as C-terminal tail (residues 402-589) revealed that CK1 bound to wild-type LIN-42 protein. Deletion of either the SYQ/CK1BD-A or LT/CK1BD-B motifs reduced this interaction (Fig. 3B).

To test whether the CK1BD is sufficient to bind CK1, we purified a truncated fragment of LIN-42 lacking the C-terminal tail (CK1BD) and performed bio-layer interferometry (BLI) assays with biotinylated CK1 and titrating LIN-42 fragments. This optical technique measures macromolecular interactions in real-time, providing *k_on_* and *k_off_* rates from which affinity constants (*K_D_*) can be calculated. The smaller size of the CK1BD compared to the CK1BD+Tail (residues 402-589; same construct used in Fig. 3B) resulted in a lower response signal, however, both proteins bound to biotinylated CK1 with comparable, nanomolar affinities (348 ± 123 nM for CK1BD and 209 ± 78 nM for CK1BD + Tail; Fig. 3C-E). These results demonstrate that the SYQ and LT motifs encompass a minimal CK1BD.

We next performed BLI experiments to quantitatively explore the significance of the CK1BD-A and -B motifs in binding to CK1. LIN-42-binding to CK1 was not detectable when CK1BD-B was deleted, either singly (*CK1BD-ΔB*) or in combination with CK1BD-A (*CK1BD-ΔA/B*) even at higher concentrations (Fig. 3C,F,G). In contrast, deletion of the CK1BD-A domain reduced the affinity of LIN-42 for CK1 approximately 10-fold (*K*_D_ = ∼3 µM)(Fig. 3C,H), necessitating higher concentrations of the CK1BD-*Δ*A +Tail protein to determine an accurate *K*_D_ (Fig. 3H). These data indicate that CK1BD-A and B both contribute to high-affinity binding of CK1 *in vitro*, while revealing a more important contribution of the CK1BD-B motif, consistent with recent work that demonstrated an essential role of the human PER2 CK1BD-B motif for stable association with CK1 in a mouse model and mammalian cells.^46^ Collectively, our data establish that the LIN-42 CK1BD is functionally conserved and mediates stable binding to the CK1 kinase domain.

### Phosphorylation of the LIN-42 C-terminus by CK1 exhibits a conserved mode of feedback inhibition

Mammalian CK1 regulates PER abundance by controlling its degradation post-translationally. Mutations on CK1 or its PER phosphorylation sites can induce changes in circadian period as great as ∼4-hours.^16,18,47,48^ To test whether LIN-42 is a CK1 substrate, we performed *in vitro* ^32^P-ATP kinase assays using the CK1BD alone or the CK1BD+Tail as substrates. CK1BD protein contains 4 serine and 6 threonine residues while the CK1BD + Tail construct contains an additional 22 serine and 15 threonine residues. We observed phosphorylation for both constructs, yet more robustly on LIN-42 CK1BD+Tail than on CK1BD alone (Fig. 4A-C). Hence, CK1 appears capable of phosphorylating both CK1BD and the extended C-terminal tail. Deletion of CK1BD-B reduced phosphorylation relative to the wild-type protein, whereas phosphorylation of the CK1BD-ΔA + Tail mutant protein remained largely unaltered (Fig. 4A-C). Thus, in the two assays, loss of CK1BD-B impacts both binding and phosphorylation by CK1 more strongly than loss of CK1BD-A (Fig. 3C, 4A-C).

**Figure 4.**
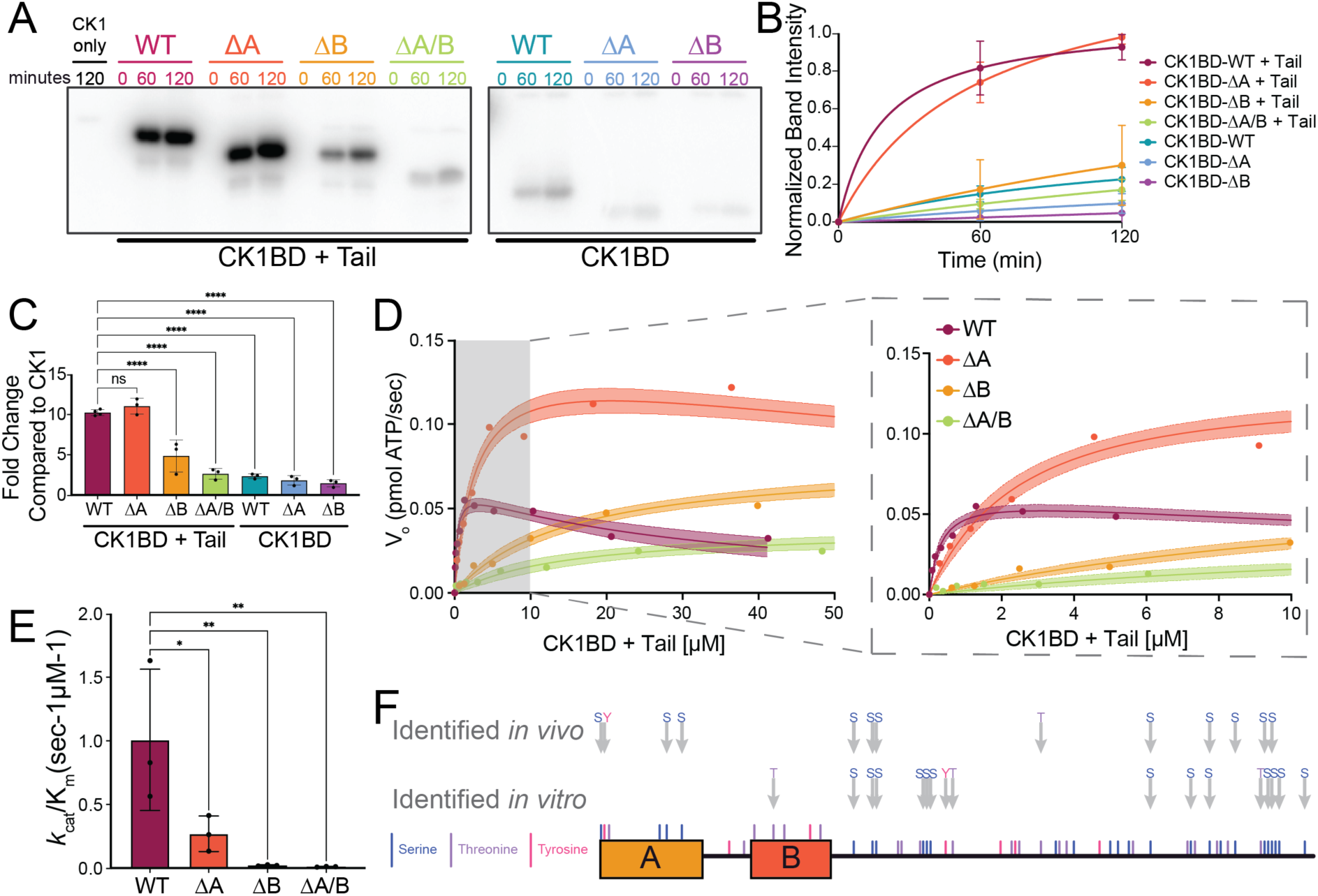
The extended C-terminus of LIN-42 is phosphorylated by CK1. **(A)** Representative ^32^P-radiolabeled ATP enzymatic assay of CK1 activity on LIN-42 CK1BD + Tail and CK1BD wild-type and indicated domain deletion mutants. **(B)** Quantification of band intensity depicted in (A) and replicates. Model fit to Michaelis-Menten, n = 3. **(C)** Fold-change quantification of band intensity from panels A & B compared to CK1 autophosphorylation at 120 minutes. Ordinary one-way ANOVA. p<0.05 was considered statistically significant. ****p < 0.0001, n ≥ 3, mean ± SEM. (D) Representative ADP-Glo enzymatic assay comparing activity of CK1 on LIN-42 CK1BD + Tail wild-type (maroon), CK1BD-ΔA (orange), CK1BD-ΔB (yellow), and CK1BD-ΔA/B (green). **(E)** Values of *k*_cat_/K_m_ calculated from (D). n=3, mean ± SEM, Ordinary one-way ANOVA. p<0.05 was considered statistically significant. * indicates p<0.03 and ** indicates p<0.002. **(F)** LIN-42 phosphorylation sites identified via mass spectrometry from *in vivo* samples from whole worm lysate collected hourly from synchronized L1 animals from 14 hrs (early L2) to 25 hrs (late L3) after plating at 25°C (top arrows) and *in vitro* kinase reactions of CK1 activity on LIN-42 CK1BD + Tail (bottom arrows). Serine, threonine, and tyrosine residues are indicated via vertical blue, purple, and pink ticks, respectively, along the CK1BD + Tail construct schematic.

To explore the kinetics of CK1-mediated phosphorylation of the LIN-42 extended C-terminus, we performed substrate titration experiments using an ADP-Glo enzymatic assay. Here, we observed a decrease in kinase activity with high levels of wild-type LIN-42 CK1BD+Tail (Fig. 4D). This result mirrors the ability of phosphorylated PER as well as the apoptosis substrate p63, to inhibit the activity of CK1 (DBT in Drosophila) in both mammalian and *Drosophila* systems through product inhibition.^14^

Closer examination revealed that deletion of the LIN-42 CK1BD-A motif relieved feedback inhibition (Fig. 4D), explaining why the ^32^P-ATP data had shown little decrease in the overall phosphorylation level (Fig. 4A-C) despite having reduced affinity for the kinase (Fig. 3C). Conversely, deletion of the B motif either in isolation or with the A motif, caused a significant reduction in overall phosphorylation of CK1BD+Tail. No serine and only 4 threonine residues are removed in this deletion, so this effect was likely not due to loss of phosphosites. The catalytic efficiency (*k*_cat_/K_m_) of CK1 for each LIN-42 substrate decreased significantly with deletion of the CK1BD motifs relative to wild-type, with activity most severely impacted upon loss of the CK1BD-B motif (Fig. 4D,E). Taken together, these results suggest that while both motifs are involved in binding to CK1, deletion of the CK1BD-A region predominantly relieves feedback inhibition without fully disrupting the anchoring interaction seen with the deletion of the CK1BD-B domain. Moreover, and similar to the mammalian PER–CK1 complex^49^, loss of the anchoring interaction compromises, but does not fully abrogate phosphorylation.

### Identification of potential LIN-42 phosphosites

To determine potential CK1-dependent LIN-42 phosphorylation sites, we performed *in vitro* kinase reactions followed by phosphoenrichment and mass spectrometry to identify phosphopeptides.There are a total of 26 serine and 21 threonine residues in the CK1BD + Tail construct. Of these, 12 serine and 3 threonine sites in the C-terminus were phosphorylated upon incubation of LIN-42 CK1BD + Tail with CK1 at least once from three replicates *in vitro* (Fig. 4F). Although we detected low levels of phosphorylation on LIN-42 constructs lacking the tail in the ^32^P-ATP assay (Fig. 4A), no serine and only 1 threonine phosphorylation within the CK1BD was detected *in vitro,* reflecting possible limitations in the assays.

To test whether LIN-42 was phosphorylated *in vivo*, we immunoprecipitated endogenously tagged 3xFLAG::LIN-42b+c at various time points from a population of synchronized L2 stage larvae and subjected precipitates to mass spectrometry. We identified 15 serine residues, 1 threonine residue, and 1 tyrosine residue that were phosphorylated on LIN-42. All but four of these phosphosites (all serine residues) are located on the CK1BD + Tail (Fig. 4F). Among these, 6 serine residues overlap with the 13 phosphoserines identified after the *in vitro* reaction while no threonine or tyrosine residues were phosphorylated in both the *in vitro* and *in vivo* datasets (Fig. 4F). The overlap of *in vitro* and *in vivo* results is consistent with the possibility that LIN-42 is a CK1 substrate *in vivo*. Whether incomplete overlap reflects differences in kinase activities in the activity of additional kinases and/or specificity factors *in vivo*, or technical differences in the experimental approaches remains to be determined.

### LIN-42 C-terminal tail deletion causes a heterochronic phenotype but leaves molting timing largely unaffected

Given the extensive phosphorylation of the LIN-42 tail *in vitro* and *in vivo*, we sought to explore its functional relevance. Unexpectedly, *lin-42(ΔTail)* mutant animals did not recapitulate the *lin-42(ΔCK1DB)* arrhythmic molting phenotype but instead resembled *lin-42(n1089)* animals phenotypically (Fig. 1C-F; Fig. 5). Specifically, *lin-42(ΔTail)* mutants exhibited regular timing of the first three molts, yet an extended and irregular fourth molt (Fig. 5A, S4A-C, S5). The severity of this extended 4^th^ molt was variable and could reflect unaccounted environmental differences during assay runs. The mutant animals also exhibited a modest reduction in brood size, egg-laying defects and precocious alae but no larval arrest (Fig. 5B-E). These data indicate that the LIN-42 C-terminus is largely dispensable for timing of larval molting and suggest instead that it plays a previously unrecognized role in the heterochronic functions of LIN-42.

**Figure 5.**
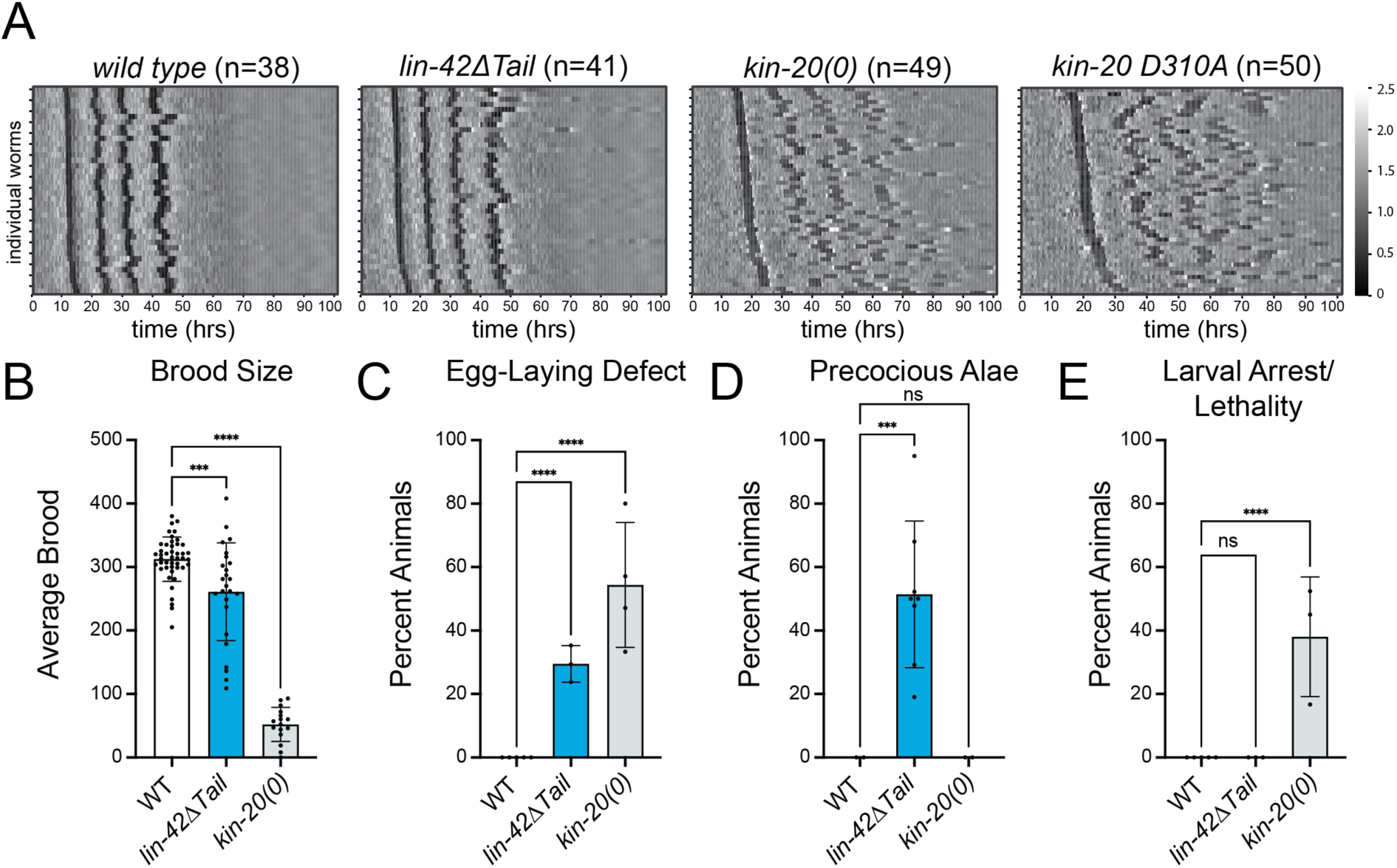
*kin-20* null and catalytically dead alleles exhibit larval arrest and asynchronous molting similar to *lin-42(ΔCK1BD)* mutants. **(A)** Heatmaps showing trend-corrected luminescence traces from the indicated genotype. Each horizontal line represents one animal. Traces are sorted by entry into the first molt. Darker color indicates low luminescence signal and corresponds to the molts. **(B)** Bar plot depicting the average number of live progeny from hermaphrodites of the indicated genotype. **(C)** Bar plot quantifying the percentage of animals of the indicated genotype that exhibited egg-laying defects as determined by the presence of hatched larvae in the animal. **(D)** Bar plot quantifying the percentage of animals of the indicated genotype with precocious complete or partial alae at the L3-L4 molt. **(E)** Bar plot quantifying the percentage of animals of the indicated genotype that arrested or died as larvae. (**B-E)** Statistical significance was determined using ordinary one-way ANOVA. p<0.05 was considered statistically significant. ** indicates p<0.0001****, indicates p<0.0001. Error bars in B-E represent standard deviation.

### *kin-20* null and catalytically dead mutants exhibit arrhythmic molting

Deletion of a major phosphorylation target, the LIN-42 tail, did not recapitulate *lin-42(ΔCK1BD)* mutant molting timing defects which prompted the question whether KIN-20 was at all required for rhythmic molting. To address this question, we monitored molt timing for *kin-20(ok505)* null mutant animals (*kin-20(0))*. *kin-20* loss recapitulated both the slow development and arrhythmic molt phenotypes of *lin-42(ok2835)* and *lin-42(ΔCK1DB)* mutations (Fig. 1B; Fig. 5A, S4A-C). Like *lin-42(ok2835)* and *lin-42(ΔCK1DB)* mutations, the *kin-20(0)* mutation also caused larval arrest when animals were grown on plate but not in the liquid culture luciferase assay (Fig. 1F; Fig. 5E). Additionally, we found that *kin-20(0)* mutant animals resembled *lin-42(ok2835)* mutants in their severe egg-laying defects and brood size reduction, phenotypes which are weaker or even absent from *lin-42(ΔCK1DB)* animals (Fig. 1D,E; Fig. 5B,C). Finally, like *lin-42(ΔCK1DB)* animals but unlike *lin-42(ok2385)* mutants, *kin-20* null mutants did not develop precocious alae (Fig. 1C; Fig. 5D), consistent with earlier work.^26^

To assess the relevance of KIN-20’s enzymatic activity for rhythmic molting, we engineered a D310A point mutation into the endogenous *kin-20* locus to abrogate catalytic activity. This mutation disrupts the HRD motif in the catalytic loop that coordinates phosphorylated residues (Fig. S3).^50,51^ The resulting *kin-20(xe355[D310A])* animals recapitulated the *kin-20(0)* mutant arrhythmic molting phenotype, although KIN-20 levels were not decreased (Fig. S6A-D). Indeed, we observed a trend towards increased accumulation of the mutant KIN-20 protein, possibly indicating autoregulation but not further pursued by us (Fig. 5A; S4A-C; S6B). We conclude that KIN-20 and especially its enzymatic activity are required for rhythmic molting.

### KIN-20 exhibits dynamic changes in subcellular localization

Although KIN-20 kinase activity and its binding to the LIN-42 CK1BD are required for rhythmic molting, LIN-42 tail phosphorylation appeared dispensable. Moreover, *kin-20* deletion or inactivation appeared to cause more pronounced defects than deletion of the LIN-42-CK1BD. Hence, we wondered whether the significance of the interaction between the two proteins could lie in regulation of KIN-20 by LIN-42, rather than vice versa. We therefore examined KIN-20 levels and localization. Using GFP::KIN-20 (with a split-GFP system to enhance the signal; Methods), we observed both nuclear and cytoplasmic signal in the epidermis. Strikingly, the relative distribution between these compartments varied with developmental stage. Microfluidics-based observation of early larvae^52^ revealed substantial cytoplasmic GFP::KIN-20 signal during molts and at the beginning of larval stages but a predominantly nuclear signal in the middle of the larval stage (Fig. S7). We recapitulated these dynamics in L4 stage animals grown on plates (Fig. 6A), which additionally revealed an apparent membrane-bound pool of KIN-20 in seam cells (Fig. 6A).

**Figure 6.**
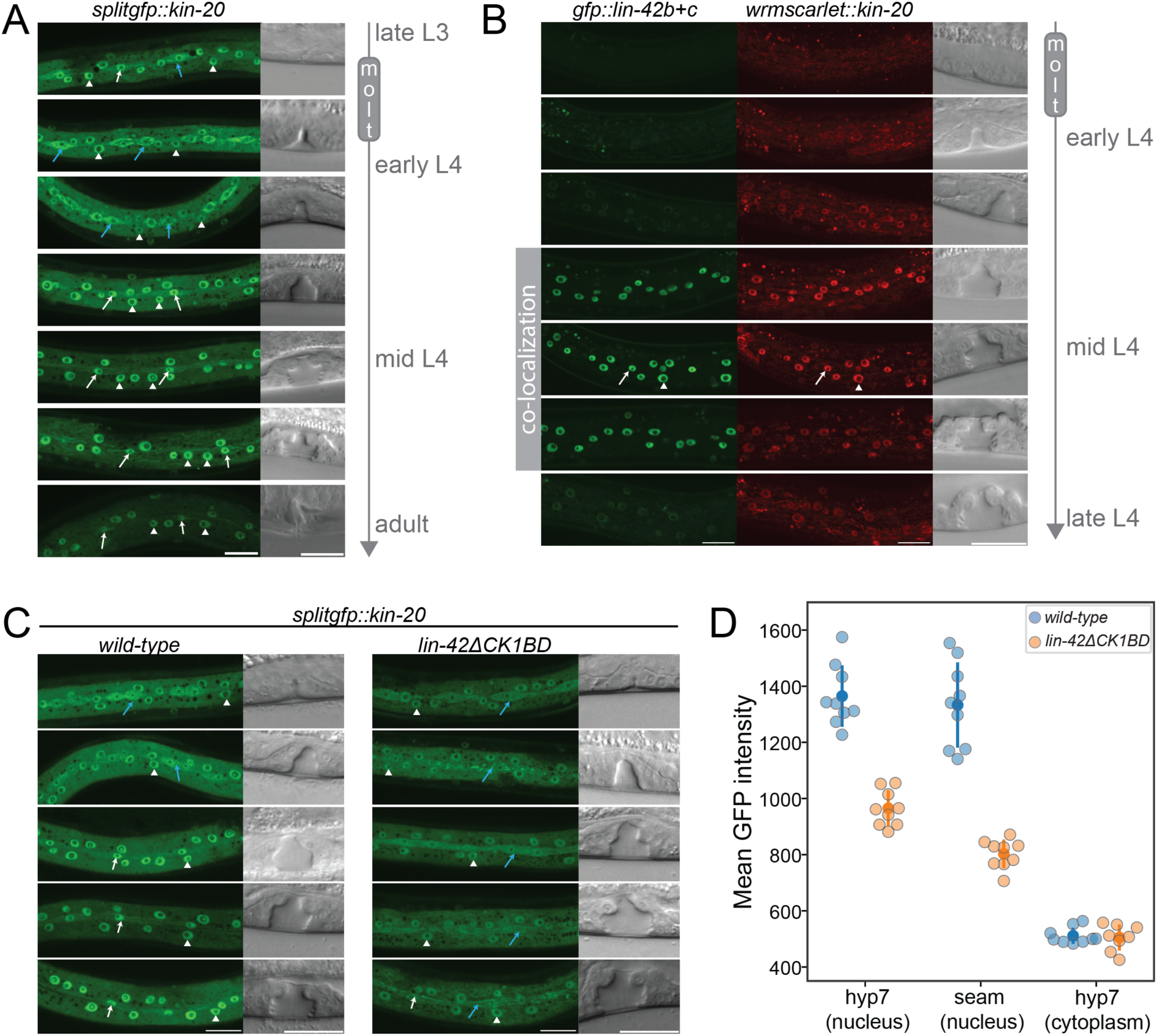
The LIN-42 CKBD is required for KIN-20 nuclear localization. **(A)** Confocal images of *splitgfp::kin-20* animals staged according to vulva morphology (Mok et al., 2015) from late L3 to adult. Arrows indicate seam cell nuclear (white) and cytoplasmic (blue) localization; white arrowheads indicate nuclear hyp7 localization. Scale bar=20 µm. **(B)** Confocal images of *gfp::lin-42b+c; wrmscarlet::kin-20* staged according to vulva morphology. An arrow indicates seam cell, an arrowhead hyp7 nuclear localization. Scale bar=20 µm. **(C)** Confocal images of *splitgfp::kin-20* in wild type and *lin-42ΔCK1BD* mutant animals during early - mid L4 stage based on the vulvae morphology. Arrows indicate seam cell nuclear (white) and cytoplasmic (blue) localization, white arrowheads indicate hyp7 nuclear localization. Scale bar=20 µm. **(D)** Quantification of GFP::KIN-20 signal in hyp7 (nucleus), seam cells (nucleus) and hyp7 (cytoplasm) of wild-type (blue) and *lin-42ΔCK1BD* (orange) animals. Each condition with n=9 mid-L4 stage worms.

### LIN-42 binding promotes nuclear localization of KIN-20

The timing of increased nuclear KIN-20 in the middle of a larval stage coincides with peak accumulation of LIN-42, especially LIN-42b (Fig. 2C). Moreover, using endogenously tagged LIN-42::GFP and wrmScarlet::KIN-20, we found that the two proteins co-localized in the nuclei of epidermal seam and hyp7 cells (Fig. 6B). Hence, we wondered if LIN-42 contributed to the change in KIN-20 localization. To test this notion, we examined how lack of LIN-42 binding affected KIN-20 localization dynamics. Specifically, we visualized GFP::KIN-20 in *lin-42(ΔCK1BD)* mutant animals during mid-L4 stage. We observed reduced nuclear GFP signal relative to wild-type animals (Fig. 6C,D) in both hypodermal and seam cell nuclei with no significant change in cytoplasmic levels. Together, these data indicate that LIN-42 promotes nuclear accumulation of KIN-20.

## Discussion

PER and its nematode orthologue LIN-42 control two distinct types of biological timing: mammalian and fly circadian rhythms (PER) and *C. elegans* developmental progression (LIN-42). The identification of additional orthologous pairs functioning in these respective pathways, ROR/NHR-23, CK1/DBT/KIN-20, Timeless/TIM-1, and REV-ERB/NHR-85, suggests that there is a conserved set of biological timing genes, whose function can be exploited in different contexts. The wiring among these components, in the few cases where this has been explored, appeared strikingly different, making it unclear why and how these genes would be evolutionarily selected for distinct timing functions. Our work shows the first example of a conserved module in these clocks: the LIN-42/PER – KIN-20/CK1 interaction.

### The PAS domains are largely dispensable for LIN-42 developmental timing function

The PAS domains are the most notable feature of LIN-42 sequence conservation and were thought to be an interaction platform for a developmental stage-specific binding partner.^19^ Previous work characterizing the *lin-42(ok2385)* and *n1089* alleles suggested that the PAS domains were necessary for the heterochronic functions of LIN-42.^22,35^ Yet, our precise deletion of the LIN-42 PAS domains caused only a mild precocious alae phenotype with significantly less penetrance than in *lin-42(n1089)* mutants, suggesting that this model is incorrect. While *lin-42(n1089)* removes the sequence of the PAS domains, it also results in a truncation of *lin-42b* and removal of part of the *lin-42a* promoter sequence.^22,35^ The *n1089* phenotype could reflect loss of the PAS domains combined with potential reduction in expression of other elements such as the C-terminal tail or an NHR-85 interaction site.^37^

The weak phenotypes seen with our precise deletion are particularly striking when considering that the PAS domains, and specifically the highly conserved PAS-B domain, mediate LIN-42 dimerization^20^ - a function that is important for PER activity in circadian rhythms^53^. Although it remains to be determined whether the PAS domains are necessary for LIN-42 dimerization *in vivo*, it seems possible that LIN-42’s molting timing function does not require dimerization.

### LIN-42 contains a CK1BD that is important for rhythmic molting

Contrasting with the PAS domain, we found the LIN-42 SYQ/LT regions to be required for rhythmic molting, and we could demonstrate *in vitro* that they constitute a functional CK1BD that anchors KIN-20/CK1, promotes LIN-42 phosphorylation, and mediates kinase inhibition. Moreover, the patterns of phosphorylation that CK1 generates on LIN-42 *in vitro* overlap with those that we observed on LIN-42 *in vivo*. We note that it remains to be demonstrated to which extent the *in vivo* phosphorylation events depend on KIN-20. Our attempts to address this question by examining LIN-42 phosphorylation in strains lacking active KIN-20 were thwarted by the slow growth, sickness, and arrhythmic molting observed in these strains.

The two CK1BD subdomains have separable functions *in vitro*. The CK1BD-B plays a more prominent role in kinase binding, mirroring the anchoring function of its mammalian and *Drosophila* equivalents.^14,46,49^ Anchoring allows CK1 to target lower affinity, non-consensus motifs, which are key regulatory sites in fly/mammalian circadian clocks.^14,46,49,54,55^ However, without the PER CK1BD-B not all capacity for phosphorylation is lost, which suggests that high-affinity binding is not essential for phosphorylation but promotes it.^49^ For LIN-42-ΔCK1BD-B, we also see low levels of phosphorylation by CK1 *in vitro* (Fig. 4), suggesting that it may play a similar role in controlling KIN-20 activity.

The LIN-42 CK1BD-A plays a less prominent role in binding but rather is involved in CK1 inhibition. CK1δ-dependent phosphorylation of both mammalian and *Drosophila* PER at sites near the CK1BD leads to feedback inhibition of CK1 through conserved anion binding sites.^14^ Our *in vitro* LIN-42 phosphorylation data reveal a decrease in enzyme catalytic activity as substrate concentration increases, dependent on the CK1BD-A, consistent with feedback product inhibition of KIN-20 (Fig. 4). Alternatively, this result could reflect substrate inhibition, where increasing amounts of LIN-42 CK1BD+Tail result in a non-productive enzyme-substrate complex. Such substrate inhibition has also been reported in several recent studies of CK1 showing phosphorylated substrates binding the enzyme and regulating its activity.^43,56–58^ Further enzymatic and structural studies will be required to distinguish between these modes of regulation.

### Function of the LIN-42-CK1 interaction

Many models of circadian clock function highlight phosphorylation of PER by CK1 as the central consequence of their interaction, controlling PER stability and thereby setting the period length. Intriguingly, although CK1 heavily phosphorylates the LIN-42 C-terminal tail, including on serine and threonine residues that are conserved among *Caenorhabditis* nematodes, but not on more distantly related nematodes (Fig. S8), deletion of this sequence does not affect molting rhythmicity. These data suggest either phosphosite redundancy, or that regulated phosphorylation of LIN-42 is not the key function of this complex for controlling molting timing.

For PER–CK1, several lines of evidence show that reciprocal to CK1-mediated regulation of PER, PER also regulates CK1 activity, both on itself and potentially other targets such as CLOCK. Indeed, PER and CRY function as a bridging complex to provide CK1 access to CLOCK^8,9^, and mutation of the CK1BD causes defects in PER and CK1 nuclear localization as well as CK1-mediated phosphorylation of CLOCK.^9,59^ Product inhibition through phosphorylated PER may thus limit phosphorylation not only of PER itself, but potentially also of other substrates, including CLOCK. Strikingly, KIN-20 also exhibits dynamic subcellular localization, with nuclear accumulation depending on LIN-42 binding. Whether this nuclear accumulation reflects co-transport, nuclear retention, or stabilization of the nuclear KIN-20 pool is not known. However, together with the fact that the LIN-42 tail, despite being heavily phosphorylated, is dispensable for rhythmic molting, this localization data indeed supports a model where a key function of the LIN-42–KIN-20 complex in the context of molting could be regulation of KIN-20 rather than LIN-42. Such a scenario may also explain the previous observation that forced expression of the short LIN-42a isoform, largely comprising the CK1BD + Tail, caused extended molt durations^35^, as this might sequester KIN-20 and/or inhibit its kinase activity.

### Distinct functions of LIN-42 and KIN-20 in molting versus heterochronic timing?

Our data not only support the existence of LIN-42 and KIN-20 in a stable complex that is important for molting timing, but also suggest that LIN-42 and KIN-20 can function independently of one another in some contexts. Neither *kin-20(lf)* nor *lin-42(ΔCK1DB)* mutations cause precocious alae, a phenotype observed with several other *lin-42* mutant alleles. These findings agree with previous genetic data that suggested independent functions of *lin-42* and *kin-20* in regulating the expression of the heterochronic *let-7* microRNA.^38^ A recent study on LIN-42’s function in regulating another heterochronic miRNA, *lin-4*, indicated that physical interaction with the NHR-85 transcription factor may support LIN-42’s activity in this process, and perhaps the heterochronic pathway more generally.^37^ Relatively mild effects upon *nhr-85* deletion, and LIN-42’s ability to interact with additional transcription factors in a yeast two-hybrid assay suggest that additional functionally relevant interactions partners remain to be identified. The apparent differences in wiring among components of the heterochronic pathway and the molting timer may reflect their different modes of timekeeping where the heterochronic pathway is chiefly concerned with the order of events and the molting timer - similar to the circadian clock - with the tempo and/or robustness of their execution.

Indeed, although LIN-42/PER have previously been suggested to provide an evolutionary link between circadian (PER) and heterochronic (LIN-42) timing systems^19^, our data suggest that the link primarily derives from functional conservation between circadian and molting timing. Hence, we may speculate that the heterochronic function of LIN-42, which appears to involve neither rhythmic activity nor conserved interactions with orthologues of other circadian clock ^19,21,23,26–28^ arose secondarily to, or independent from, a rhythmic timing function. Finally, we note with interest that the phosphorylation of PER by CK1 is an important component of temperature compensation in the circadian clock^60^, which allows this clock to maintain its period despite variation in temperature. By contrast, the period of *C. elegans* molting and oscillatory genes expression changes with temperature. Our data that the LIN-42 tail, where the bulk of CK1 mediated phosphorylation occurs, is dispensable for rhythmic molting may thus reflect the lack of robust temperature compensation in the *C. elegans* developmental clock.

## Materials and Methods

### C. elegans culture

*C. elegans* were grown and maintained at 20°C unless indicated otherwise and cultured as described^61^ on MYOB^62^ or NGM 2% agar plates with *Escherichia coli* OP50 bacteria. Full genotypes of strains used in this work and the respective growth conditions are listed in Table S2. All mutant strains were backcrossed to N2 at least 2X. The Bristol N2 isolate was used as the wildtype. Below, we describe how we generated novel strains for this study. In Table S3, we list oligo repair templates, plasmids and genotyping primers. In Table S4 we list crRNAs used.

### Genome editing

Novel *lin-42* mutant alleles (*(wrd67[ΔPAS])*, (*wrd63[ΔCK1BD)* and (*wrd107[ΔTail])*) using CRISPR-Cas9 were generated by injection of Cas9 or Cas12 ribonucleoprotein complexes as described.^63^ Specified deletions were made in the endogenous *lin-42* locus in N2 animals. Repair templates were used at 100 ng/µl along with Cas9 or Cas12 at 250 ng/µl, crRNA oligos at 60 ng/µl, and 10 ng/µl co-injection marker (pCFJ90).^64^ Strains used in the luciferase assays were generated by crossing the mutant strains with a previously described luciferase reporter strain (HW1993)^40^ (Table S2).

Strains containing endogenously tagged *lin-42* and *kin-20* were obtained using CRISPR/Cas9 as described previously.^64^ In short: 5 µg of of Alt-R S.p. Cas9 Nuclease V3 (IDT, Cat # 1081058), 2 µg of Alt-R® CRISPR-Cas9 tracrRNA (IDT, Cat # 1072532) and 1.5 µg of crRNA were incubated at 37°C for 15 min to form the RNP complex. 500 ng repair template and co-injection markers pIK127 (10 ng/μl) and pRF4 (*rol-6(su1006)*) (40 ng/μl) were added to a final volume of 20 μl in water. Mix was injected into N2 animals. For *lin-42*, a *gfp::3xflag* (amplified from plasmid pIK384 with primers KK41/KK42) or *3xflag* tag (Ultramer DNA Oligo, KK90) was inserted at the N-terminus of LIN-42b+c isoforms.

*For kin-20*, a *3x::flag-ha* (Ultramer DNA oligo, KK107), *4xgfp11::linker:.flag* (amplified from plasmid pIK401 with primers KK145/KK146) or *wrmScarlet* (amplified from plasmid pIK385 with primers KK102/KK103) was inserted at the first common shared exon of all KIN-20 isoforms with following flanking sequence: 5′ tatattcaaattttcagCGGAGATG-[insert]*-*GAACTTCGTGTCGGCAATCGTTTCC 3′).

To reconstitute GFP with the split-GFP system, we crossed the obtained *4xgfp11::linker::flag::kin-20* line into a MosSCI strain containing the *gfp1-10* fragment expressed from an *eft-3* promoter. For this strain the *gfp1-10* sequence from plasmid pCZGY2254^65^ for *C. elegans* was codon optimized. After adding two synthetic introns the codon-optimized sequence was ordered as a gBlock from IDT. The sequence was cloned into a MosSCI-compatible backbone together with *eft-3*p promoter and *tbb-2* 3’UTR sequences to create plasmid pIK407 by Gibson assembly. pIK407 (*eft-3p::gfp1-10(codon-optimized)::tbb-2* 3’UTR) was inserted into the ttTi5605 site on chromosome II by MosSCI.^66^

To generate a catalytic dead KIN-20, the D310A mutation was introduced by CRISPR-Cas9 into N2 and *3xflag-ha::kin-20* animals, respectively. Injection was performed as described previously for the *lin-42* and *kin-20* tagged strains. The ultramer DNA oligo KK137 containing the D310A mutation and two silent mutations (to abrogate crRNA recognition) was used as a repair template.

### Immunoprecipitation

For the 3xFLAG-LIN-42 IP, mixed stage animals were obtained by plating eggs after a standard hypochlorite treatment and growth at different temperatures (15°C, 20°C and 25°C). 24 hrs after plating, worms were collected by pooling 30,000 worms from each temperature condition per strain and replicate (triplicates for each strain). For the HA-KIN-20 IP, synchronized worms were collected after 32 hrs at 25°C. For the mapping of LIN-42 phosphorylation sites, synchronized worms were grown at 25°C and collected hourly from 14 hrs (L2 larval stage) to 25 hrs (L4 larval stage). Worm extracts followed by immunoprecipitation were done as previously described.^67^ In brief, extracts were made in lysis buffer (50mM Tris-HCl, pH 7.5, 150 mM NaCl, 1% TRITON X-100, 1 mM EDTA) supplemented with 1 mM PMSF and 1 tablet of cOmplete Protease inhibitor (Roche, cat. No. 11 873 580 001) per 50 ml using an MP Biomedical Fast-Prep-24 5G bead beater. For the FLAG IP, cleared worm extract was incubated with 50 μl of 50% bead suspension of anti-FLAG M2 Magnetic Beads (Sigma. Catalog Number M8823) for 2 hours at 4°C. For the HA-IP worm extract was incubated with 50 µl of bead suspension of Pierce™ Anti-HA Magnetic Beads (Thermo Scientific, Cat. No. 88837).

### Proteolytic digest

The beads were incubated for 4 hrs at RT with 800 rpm shaking in 5 μl of digestion buffer (3 M guanidine hydrochloride, 20 mM EPPS (4-(2-Hydroxyethyl)-1-piperazinepropanesulfonic acid) pH 8.5, 10 mM CAA (2-Chloroacetamide), 5 mM TCEP (Tris(2-carboxyethyl)phosphine hydrochloride)) and 1 μl of Lys-C (0.2 μg/μl in 50 mM^68^, pH 8.5). 1 μl of trypsin (0.2 μg/μl) was added and incubated at 37°C overnight. After 12 hrs another 1 μl of trypsin was added and incubation continued for 4 more hours at 37°C

### Mass spectrometry to identify LIN-42 interacting factors

The generated peptides were acidified with 0.8% TFA (final concentration) and analyzed by LC–MS/MS on an EASY-nLC 1000 [Thermo Scientific] with a two-column set-up. The peptides were applied onto a peptide 75 µm x 2 cm PepMap Trap trapping column [Thermo Scientific] in 0.1% formic acid, 2% acetonitrile in H2O at a constant pressure of 800 bar. Using a flow rate of 150 nl/min, peptides were separated on a 50 µm x 15 cm PepMap C18, 2 µm, 100 A [Thermo Scientific] at 45 °C with a linear gradient of 2%–6% buffer B in buffer A in 3 min followed by a linear increase from 6 to 22% in 40 min, 22%– 28% in 9 min, 28%–36% in 8 min, 36%–80% in 1 min, and the column was finally washed for 14 min at 80% buffer B in buffer A (buffer A: 0.1% formic acid; buffer B: 0.1% formic acid in acetonitrile). The separation column was mounted on an EASY-Spray™ source [Thermo Scientific] connected to an Orbitrap Fusion LUMOS [Thermo Scientific]. The data were acquired using 120,000 resolution for the peptide measurements in the Orbitrap and a top T (3 s) method with HCD fragmentation for each precursor and fragment measurement in the ion trap according the recommendation of the manufacturer (Thermo Scientific). For the analysis protein identification and relative quantification of the proteins was performed with MaxQuant v.1.5.3.8 using Andromeda as search engine, and label-free quantification (LFQ). The *C. elegans* subset of the UniProt v.2021_05 combined with the contaminant database from MaxQuant was searched and the protein and peptide FDR were set to 0.01. The LFQ intensities estimated by MaxQuant were analyzed with the einprot R package (https://github.com/fmicompbio/einprot) v0.5.4. Features classified by MaxQuant as potential contaminants or reverse (decoy) hits or identified only by site, as well as features identified based on a single peptide or with a score below 10, were filtered out. The LFQ intensities were log2 transformed and missing values were imputed using the ‘MinProb’ method from the imputeLCMD R package v2.0 with default settings. Pairwise comparisons were performed using limma v3.50.0, considering only features with at least 2 non-imputed values across all the samples in the comparison. Estimated log2-fold changes and P-values (moderated t-test) from limma were used to construct volcano plots. Data will be available via ProteomeXchange with identifier PXD058601 upon publication.

### Mass spectrometry to map *in vivo* LIN-42 phosphorylation sites

The peptides from each sample were labeled with TMTpro reagents and pooled. The TMT labeled peptide mixture was subjected to off-line high pH fractionation on a YMC Triart C18 0.5 × 250 mm column (YMC Europe GmbH) using the Agilent 1100 system (Agilent Technologies). A total of 96 fractions was collected for each experiment and concatenated into 48 fractions as previously described. For each LC-MS analysis, all available peptides were loaded onto a PepMap Neo trap (Thermo Fisher) using the Vanquish Neo UHPLC system (Thermo Fisher). On-line peptide separation was performed on a 15-cm EASY-Spray™ C18 column (ES75150PN, Thermo Fisher) by applying a linear gradient of increasing ACN concentration at a flowrate of 200 nL/min. Orbitrap Fusion Lumos Tribrid (Thermo Fisher) mass spectrometer was operated in the data-dependent mode. The ions for the survey scan were collected for a maximum of 50 ms to reach the standard AGC target value and the scan recorded using an Orbitrap detector at a resolution of 120,000. The topmost intense precursor ions from the Orbitrap survey scan recorded every 3 sec were selected for stepped higher-energy C-trap dissociation (HCD) at 29%, 32% and 35% normalized collision energy scan. To reach an AGC value of 100,000 ions, the maximum ion accumulation time for the MS2 scan was set to 500 ms. The TMT reporter ions were quantified using an MS2 scan recorded using the Orbitrap analyzer at a resolution of 50,000. Thermo RAW files were processed using Proteome Discoverer 2.4 software (Thermo Fisher) as described in the manufacturer’s instructions. Briefly, the Sequest search engine was used to search the MS2 spectra against the *C. elegans* UniProt database (downloaded on 05/2023) supplemented with common contaminating proteins. For peptide identification, cysteine carbamidomethylation and TMTpro tags on lysine and peptide N-termini were set as static modifications, whereas oxidation of methionine residues and acetylation protein N-termini were set as variable modifications. The assignments of the MS2 scans were filtered to allow 1% FDR. For reporter quantification, the S/N values were corrected for isotopic impurities of the TMTpro reagent using the values provided by the manufacturer. The sums across all TMTpro reporter channels were normalized assuming equal total protein content in each sample. Data will be available via ProteomeXchange with identifier PXD058598 upon publication.

### Western Blot

Synchronized animals were collected hourly from 13 hrs - 23 hrs after plating at 25°C. Extracts were made by boiling at 95°C for 5 minutes in lysis buffer (63 mM Tris-HCl (pH 6.8), 5 mM DTT, 2% SDS, 5% sucrose) followed by sonication with a BioRupter Plus (Diagnode) with the following settings: 13 cycles, 30 s on/off at 4°C. Samples were cleared by centrifugation, before separating proteins by SDS-PAGE (loading: 50 µg protein extract per well) and transferring them to PVDF membranes by semi-dry blotting. The following antibodies were used: Monoclonal mouse anti-FLAG M2-Peroxidase (HRP)(Sigma-Aldrich; A8592, dilution: 1:1000), monoclonal mouse anti-Actin clone C4 (Millipore; MAB1501, dilution 1:7500) and anti-mouse HRP conjugated antibody (GE Healthcare #NXA931, dilution 1:2000). The membrane was incubated with ECL™ Prime Western-Blot-Reagent (Cytiva, #RPN2236) and bands detected with an Amersham Imager 680 (GE Healthcare).

### Phenotypic analysis

For phenotypic analyses, gravid adults were bleached, and embryos were hatched at low density onto MYOB plates seeded with OP50, or gravid adults were picked onto seeded plates and removed after 1-2 hours. Animals were maintained at 20°C and observed daily or twice daily. Bag-of-worms phenotype was determined when live progeny were observed inside adult animals. For brood counts, animals were individually transferred to fresh wells daily for five days after L4 and live progeny were counted and averaged.

To score premature alae, synchronized animals were collected from MYOB plates by washing off plates. 1000 µl of M9 + 2% gelatin was added to the plate or well, agitated to suspend animals in M9+gelatin, and then transferred to a 1.5 ml tube. Animals were spun at 700xg for 1 min. The media was then aspirated off and animals were resuspended in 500µl M9 + 2% gelatin with 5 mM levamisole. 12 µl of animals in M9 +gel with levamisole solution were placed on slides with a 2% agarose pad and secured with a coverslip. Images were acquired using a Plan-Apochromat 40x/1.3 Oil DIC lens or a Plan-Apochromat 63x/1.4 Oil DIC lens on an AxioImager M2 microscope (Carl Zeiss Microscopy, LLC) equipped with a Colibri 7 LED light source and an Axiocam 506 mono camera. Acquired images were processed through Affinity photo software (version: 1.9.2.1035). For the confocal microscopy, L3/L4 stage-animals were picked from OP50 plates grown at 25°C. Worms were mounted on a glass slide with a 2 % agarose patch immobilized with 5 μl of 10 mM levamisole (Fluca Analytical, #31742). Images were acquired in channels for red (561 nm laser), green (488 nm laser) and DIC with a 40x/1.3 immersion objective on a Zeiss LSM700 confocal microscope. Acquired images were processed with FiJi.^68^

### Quantification of GFP levels

For quantification, mid-L4 stage worms (defined by vulval morphology) were selected. Images acquired in the GFP fluorescence channel were imported into Napari for analysis.^69^ The following regions of interest were manually annotated for quantification: Nuclei of hyp7 and seam cells. For the cytoplasm of hyp7, the layer with the maximum intensity of nuclear GFP signal in hyp7 was selected and a region between the annotated hyp7 nuclei was chosen. To define the background, a region in the gonad was used (where no GFP signal was detected). For the quantification, average GFP levels were computed per region and worm. Background levels were subtracted to the GFP levels of all other regions.

### Microfluidics

Worms were prepared and loaded into the device as described in Berger et al. (2021). Briefly, gravid adult hermaphrodites were harvested and treated with bleach. The resulting embryos were collected by centrifugation at 1,600 g for 1 minute and washed three times with S-Basal buffer. For synchronization, embryos were incubated overnight in S-Basal, allowing larvae to arrest. Arrested larvae were then passed through a 10 µm filter (pluriStrainer Mini 10 µm, PluriSelect), and 6,000 worms were transferred to 3 NGM plates, where they were incubated for 12 hours at 25°C. Once the worms reached the late L1 stage, they were harvested using M9 buffer, washed twice to remove debris, and loaded into the experimental device. During the experiment, bacterial food was supplied at a constant rate of 1 µl/h, with periodic increases to 100 µl/h for 5 seconds every 30 minutes to clear debris. Images were acquired every 10 minutes over 24 hours using a spinning disk confocal scanning microscope (Yokogawa CSU W1 with Dual T2). Brightfield and fluorescent signals (488 nm laser) were recorded simultaneously using two sCMOS Photometrics Prime 95B cameras with a 40 x oil immersion lens (NA = 1.3). Imaging was conducted with a 25 ms exposure time and a motorized z-drive, acquiring z-stacks with a 1 µm step size and 20 images per stack. Analysis was performed using Fiji/ImageJ software.

### Luciferase assays

Assays were performed and analyzed as described.^40^ Briefly, gravid adults were bleached, and eggs were immediately singled into wells containing 90 µl OP50/S-Basal/D-Luciferin solution per well and left in the luminometer machine and measured every 10 minutes for 0.5 seconds. The assays lasted 96 hrs or 130 hrs. For statistical analysis, we performed a Wilcoxon-Mann-Whitney test (implemented in the python package SciPy version 1.4.1 as the function Mann-Whitney-U).

### Expression and purification of recombinant proteins

All proteins were expressed from a pET22-based vector in *Escherichia coli* Rosetta (DE3) cells based on the Parallel vector series.^70^ All LIN-42 constructs (CK1BD + Tail, residues 402-598; CK1BD, residues 402-475; CK1BD-ΔA, -ΔB, and ΔAB mutants) were expressed downstream of an N-terminal TEV-cleavable His-NusA tag. Human CK1δ catalytic domains (CK1δ ΔC, residues 1–317) were all expressed with a TEV-cleavable His-GST tag. All proteins expressed from Parallel vectors have an additional N-terminal vector artifact of ‘GAMDPEF’ remaining after TEV cleavage. Cells were grown in LB media at 37°C until the O.D.600 reached ∼0.8; expression was induced with 0.5 mM IPTG, and cultures were grown for approximately 16–20 hr more at 18°C. Cells were centrifuged at 3,200 x g, resuspended in 50 mM Tris, pH 7.5, 300 mM NaCl, 20 mM imidazole, 5% (vol/vol) glycerol, 1 mM tris(2-carboxyethyl)phosphine (TCEP), and 0.05% Tween-20. For purification of recombinant protein, cells were lysed with a microfluidizer followed by sonication and then the lysate was clarified via centrifugation at 140,500 x g for 1 hour at 4°C. Ni-NTA affinity chromatography was used to extract his-tagged proteins from the lysate and then the affinity/solubility tags were cleaved using His6-TEV (GST-TEV for CK1δ ΔC) protease overnight at 4°C. The cleaved protein was then separated from solubility tag and TEV by a second Ni-NTA (GST for CK1δ ΔC) affinity column and further purified using size exclusion chromatography (SEC) in 50 mM Tris, pH 7.5, 200 mM NaCl, 1 mM EDTA, 5% (vol/vol) glycerol, 1 mM TCEP, and 0.05% Tween-20. Small aliquots of protein were frozen in liquid nitrogen and stored at −70°C for long-term storage.

### *In vitro* biotinylation, pull-down assays, and bio-layer interferometry

LIN-42 CK1BD + Tail constructs and CK1δ ΔC were biotinylated via Sortase A-mediated reactions between a Sortase A recognition motif peptide (biotin-LPETGG) and our LIN-42 CK1BD + Tail/CK1δ ΔC (N-terminal G from ‘GAMDPEF’ artifact). Reactions were carried out in 50 mM Tris pH 7.5 and 150 mM NaCl using 5 µM His6-Sortase A, 300 µM biotin-LPETGG, and 50 µM protein. Ni-NTA affinity chromatography followed by SEC was used to purify labeled protein from His6-Sortase A and excess biotin, respectively. Pull down assays were performed using magnetic streptavidin beads to bind biotinylated LIN-42 CK1BD + Tail WT and mutants (final concentration 5 µM) in the presence and absence of CK1δ ΔC (final concentration 5 µM). All BLI assays were performed in SEC buffer supplemented with 7.5mM BSA as previously described^71,72^ using an eight-channel Octet-RED96e.

### ADP-Glo kinase assays

Substrate titration kinase reactions were performed on the indicated recombinant LIN-42 proteins (CK1BD + Tail WT or mutants) using the ADP-Glo kinase assay kit (Promega) according to the manufacturer’s instructions. All reactions were performed in 30 µL volumes in duplicate (n=3 independent experiments) using 1x kinase buffer (25mM Tris pH 7.5, 100 mM NaCl, 10 mM MgCl_2_, and 2 mM TCEP) supplemented with 100µM ATP, 0.2µM recombinant CK1, and indicated LIN-42 proteins. Reactions were held at room temperature for 1 hour and then 5 µL aliquots were taken and quenched with ADP-Glo reagents. Luminescent measurements were measured at room temperature with a SYNERGY2 microplate reader in 384-well microplates. Data analysis was performed using Excel (Microsoft) and Prism (GraphPad).

### ^32^P-ATP kinase assays

1 µM CK1δ ΔC was incubated with 10 µM LIN-42 (CK1BD + Tail/CK1BD WT and mutant proteins) in 1x kinase buffer (25 mM Tris pH 7.5, 100 mM NaCl, 10 mM MgCl2, 2 mM TCEP). Reactions were started by the addition of ^32^P-ATP (final concentration 2 mM) and samples were collected at indicated time points and quenched in an equivalent volume of 2x SDS-PAGE loading buffer. Proteins labeled with ^32^P were separated and analyzed via SDS-PAGE and the gels were dried at 80°C for 2 hours before overnight exposure in a phosphor screen (Amersham Biosciences). A Typhoon Trio (Amersham Biosciences) phosphorimager was used to visualize exposed gels and ^32^P-labeled protein bands were quantified via densitometry using ImageJ (NIH), Excel (Microsoft) and Prism (GraphPad).

### *In vitro* Mass Spectrometry

#### Sample preparation

Kinase reactions were performed in 1x kinase buffer (25 mM Tris pH 7.5, 100 mM NaCl, 10 mM MgCl2, 2 mM TCEP) for 60 minutes and then quenched with 20mM EDTA. Samples were denatured, reduced, alkylated, and digested according to the In-solution Protein Digestion (Promega) protocol. In brief, phosphorylated samples were denatured/reduced in 8M urea/50mM ammonium bicarbonate/5mM Dithiothreitol (DTT) for 1hr at 37°C. Samples were incubated for 30 minutes in the dark with 15mM iodoacetamide and then digested with X Grade Modified Trypsin according to the Trypsin Digestion Protocol (Promega). Digested Samples were phospho-enriched using the High-Select Fe-NTA Phosphopeptide Enrichment Kit (Thermo Scientific) and sent for analysis at the University of California, Davis Proteomic Core Facility.

#### LC-MS

For each sample, equal volumes were loaded onto a disposable Evotip C18 trap column (Evosep Biosytems, Denmark) as per the manufacturer’s instructions. Briefly, Evotips were wetted with 2-propanol, equilibrated with 0.1% formic acid, and then loaded using centrifugal force at 1200g. Evotips were subsequently washed with 0.1% formic acid, and then 200 μL of 0.1% formic acid was added to each tip to prevent drying. The tipped samples were subjected to nanoLC on a Evosep One instrument (Evosep Biosystems).

Tips were eluted directly onto a PepSep analytical column, dimensions: 15 cm x 75 um C18 column (PepSep, Denmark) with 1.5 μm particle size (100 Å pores) (Bruker Daltonics), and a ZDV spray emmiter (Bruker Daltronics). Mobile phases A and B were water with 0.1% formic acid (v/v) and 80/20/0.1% ACN/water/formic acid (v/v/vol), respectively. The standard pre-set method of 40 samples-per-day whisper method was used, which is a 31-minute run.

Mass Spectrometry – Performed on a hybrid trapped ion mobility spectrometry-quadrupole time of flight mass spectrometer (timsTOF Pro, (Bruker Daltonics, Bremen, Germany) with a modified nano-electrospray ion source (CaptiveSpray, Bruker Daltonics). In the experiments described here, the mass spectrometer was operated in diaPASEF mode. Desolvated ions entered the vacuum region through the glass capillary and deflected into the TIMS tunnel which is electrically separated into two parts (dual TIMS). Here, the first region is operated as an ion accumulation trap that primarily stores all ions entering the mass spectrometer, while the second part performs trapped ion mobility analysis.

#### DIA PASEF

The dual TIMS analyzer was operated at a fixed duty cycle close to 100% using equal accumulation and ramp times of 85 ms each.

Data-independent analysis (DIA) scheme consisted of one MS scan followed by MSMS scans taken with 19 precursor windows at width of 50Th per 0.57s cycle over the mass range 300-1200 Dalton. The TIMS scans layer the doubly and triply charged peptides over a ion mobility −1/k0-range of 0.7-1.3 V*sec/cm2. The collision energy was ramped linearly as a function of the mobility from 59 eV at 1/K0=1.4 to 20 eV at 1/K0=0.6.

#### Data Analysis

##### DIA

LCMS files were processed with Spectronaut version 16.1 (Biognosys, Zurich, Switzerland) using DirectDIA analysis mode. Mass tolerance/accuracy for precursor and fragment identification was set to default settings. The unreviewed FASTA for *C. elegans* was downloaded from Uniprot and a database of common laboratory contaminants were used.^73^ A maximum of two missing cleavages were allowed, the required minimum peptide sequence length was 7 amino acids, and the peptide mass was limited to a maximum of 4600 Da. Carbamidomethylation of cysteine residues was set as a fixed modification, and methionine oxidation and acetylation of protein N termini as variable modifications. A decoy false discovery rate (FDR) at less than 1% for peptide spectrum matches and protein group identifications was used for spectra filtering (Spectronaut default). Decoy database hits, proteins identified as potential contaminants, and proteins identified exclusively by one site modification were excluded from further analysis.

#### Data availability

Data is available from the massive online repository https://massive.ucsd.edu/ ID number MSV000096641 and proteome Exchange ID number PXD058784

## Supporting information

Combined Supplemental Figures

## Author contributions

**Rebecca K. Spangler:** Conceptualization; formal analysis; investigation; writing – original draft. **Guinevere E. Ashley:** Conceptualization; formal analysis; investigation; writing – original draft. **Kathrin Braun:** Conceptualization; Formal analysis; investigation; writing – review and editing. **Marit van der Does:** Formal analysis; investigation; writing – review and editing. **Daniel Wruck:** Formal analysis; investigation; writing – review and editing. **Andrea Ramos-Coronado:** Formal analysis; investigation; writing – review and editing. **James Matthew Ragle:** Formal analysis; investigation; writing – review and editing. **Vytautas Iesmantavicius:** Formal analysis; investigation; writing – review and editing. **Lucas J Morales Moya:** Formal analysis; writing – review and editing. **Keya Jonnalagadda:** Formal analysis; writing – review and editing. **Carrie L. Partch:** Conceptualization; supervision; funding acquisition; writing – review and editing; project administration. **Helge Großhans:** Conceptualization; supervision; funding acquisition; writing – original draft; project administration. **Jordan D. Ward:** Conceptualization; supervision; funding acquisition; writing – original draft; project administration.

## Acknowledgements

Some strains were provided by the *Caenorhabditis* Genetics Center, which is funded by the NIH Office of Research Infrastructure Programs [P40 OD010440]. We thank Iskra Katic for worm injections and technical support, Daniel Hess and Jan Seebacher for mass spectrometry analysis, and Dimos Gaidatzis for help with proteomics data analysis. We are thankful to Laurent Gelman and Laure Plantard for help with confocal imaging. Some analysis for this project was performed in the Proteomics Core Facility of the Genome Center, University of California, Davis with instrument funding provided by the NIH (S10OD026918-01A1). We thank Michelle Salemi (UC Davis) for performing the proteomics sample prep, the LC-MS/MS method writing and running and data analysis. The Bruker tims-TOF HT MS/Evosep LC system was supported by the Howard Hughes Medical Institute Investigator Award for Dr. Neal Hunter, UC Davis. Biolayer Interferometry data were collected with instrument funding provided by the NIH (S10OD027012).

## Funding

This work was funded by the National Institutes of Health (NIH) National Institute of General Medical Sciences (NIGMS) to J.D.W (R01GM138701) and C.L.P. (R35GM141849), and the Howard Hughes Medical Institute to C.L.P. This work is part of a project that has received funding from the European Research Council (ERC) under the European Union’s Horizon 2020 research and innovation program (Grant agreement No. 741269, to H.G.) and from the Swiss National Science Foundation (#310030_207470, to H.G.). The FMI is core-funded by the Novartis Research Foundation.

## Data availability

All relevant data can be found within the article and its supplementary information.

---

**Figure S1 PER proteins have a crucial role in mammalian circadian rhythms. (A)** Cartoon schematic of the primary transcription-translation feedback loop which generates ∼24-hour rhythms in mammals. CCGs, Clock-controlled genes. **(B)** Cartoon schematic illustrating the mammalian phosphoswitch that dictates CK1-dependent regulation of PER stability. Crystal structure of human CK1δ (gray) bound to phosphorylated PER2 FASP (orange, 4pFASP) peptide, PDB: 8d7o. pD, phosphodegron; FASP, Familial Advanced Sleep Phase.

**Figure S2 Larval stage durations for *lin-42* mutant animals.** Boxplots showing durations (in hours) for **(A)** larval stages **(B)** molts and **(C)** intermolts from luciferase assay. Wild type in white, *lin-42(n1089)* in dark grey, *lin-42(ok2385)* in light grey, *lin-42ΔPAS* in blue and *lin-42ΔCK1BD* in green. **(D)** Bar plot showing the number of molts from the luciferase assay of the indicated genotypes.

**Figure S3 Conservation and structural prediction of *C. elegans* KIN-20. (A)** Sequence alignment of *C. elegans* KIN-20 and KIN-19, *H. sapiens* CK1δ, *M. musculus* CK1δ, *A. thalia* CK1, *S. cerevisiae* HRR25 (HRR), and *D. melanogaster* Doubletime (DBT). Important enzymatic sequence features are boxed; full kinase domain (blue box - 79% identical between KIN-20 and human CK1δ), catalytic lysine (purple - conserved in KIN-20), catalytic loop (pink - 100% conserved between KIN-20 and human CK1δ), Magnesium (Mg)-binding loop (orange - 12 out of 13 residues conserved between KIN-20 and human CK1), anion coordination sites 1 (yellow), 2 (green), and 3 (blue) (all conserved in KIN-20). **(B)** Crystal structure of *H. sapiens* CK1δ (PDB 6pxo). Dark blue indicates residue is conserved in KIN-20; residues that diverge are highlighted in pale blue (similar amino acid) or not conserved (white). **(C)** AlphaFold structural prediction of KIN-20 (https://alphafold.ebi.ac.uk/entry/A8X4B3) colored by the model confidence. **(D)** KIN-20 AlphaFold plot of predicted aligned error.

**Figure S4 Larval stage durations for *kin-20 and lin-42* mutant animals.** Boxplots showing durations (in hours) for **(A)** Larval stages **(B)** Molts **(C)** Intermolts from luciferase assay. Wild type (*Wt*) in white, *lin-42(ΔTail)* in blue, *kin-20(0)* in light grey, *kin-20 D310A* in dark grey. Statistics were done using the Mann-Whitney U-test. Stars indicate the significance of difference between the *Wt* strain and the different *lin-42* and *kin-20* mutant animals: * p<0.05, ** p<0.01, *** p<0.001, **** p<0.0001. **(D)** Bar plot showing the number of molts detected in the assay in percentage of animals.

**Figure S5 Luciferase replicate of *lin-42(ΔTail).*** Heatmaps showing trend-corrected luminescence traces from the indicated genotype. Each horizontal line represents one animal. Traces are sorted to the entry of the first molt. Darker color indicates low luminescence signal and corresponds to the molts.

**Figure S6 *kin-20(D310A)* affects molt timing. (A)** Western Blot with extracts from *3xflag::lin-42b+c, 3xflag::kin-20* and *3xflag::lin-42b+c; 3xflag::kin-20* larvae (mid L4 stage). Blot probed with anti-FLAG-HRP (1:1000). Arrows indicate bands for LIN-42b and KIN-20. **(B)** Western Blot with extracts from *3xflag::kin-20* wild-type, *(xe401[D310A])* and *(xe400[D310A])* mutant animals. Top panel probed with anti-FLAG-HRP (1:1000). Lower panel probed with anti-actin-1 (1:7500). Arrows indicate bands for KIN-20. **(C)** Heatmaps showing trend-corrected luminescence traces from the indicated genotype. Each horizontal line represents one animal. Traces are sorted by entry into the first molt. Darker color indicates low luminescence signal and corresponds to the molts. **(D)** Boxplots showing the duration (in hours) for the larval stage from the luciferase assay.

**Figure S7 KIN-20 dynamics during L2 - L3 stage.** Microscopy images of one *splitgfp::kin-20* larva from a microfluidics experiment at indicated timepoints. Time indicated in minutes after Molt 1 (M1) or Molt 2 (M2) exit. Arrows indicate seam cell nuclear (white) and cytoplasmic (blue) localization; white arrowheads indicate nuclear hyp7 localization. Scale bar=50 µm.

**Figure S8 Alignment of nematode LIN-42 protein sequences.** LIN-42 homologs from the indicated nematode species were aligned using Clustal Omega. The length in amino acids of each homolog follows the species and homolog name. To the left and right of the alignment are amino acid positions of the end residues for each protein. Blue shading indicates conserved sequences and the histogram at the bottom depicts the degree of conservation with a consensus sequence listed below. The positions of the *C. elegans* CK1BD-A and CK1BD-B motifs are indicated. The location of the phosphosites found in our *in vivo*, *in vitro* and both datasets are indicated.

## TABLES

**Table S1 *lin-42* mutant phenotypes**

All animals grew from eggs hatched onto seeded plates at low density after sodium hypochlorite treatment. n ≥ 20 for all analyses. ^a^Percentage of animals that failed to reach adulthood by day 8 after hatching or died as young larva. ^b^Percentage of animals with alae formation in early L4 (L4.0-L4.2). ^c^Percentage of animals that exhibited bag-of-worms phenotype. ^d^Average number of progeny from fertile adults; arrested and bagged animals were not included in the calculation. *Broods were not counted in animals with high instance of BOW phenotype.

**Table S2 strains**

*C. elegans* strains used in this work.

**Table S3 oligos and plasmids**

Oligos and plasmids used in this work

**Table S4 crRNAs**

CRISPR RNAs used in this work

## Notes

### Competing Interest Statement

The authors have declared no competing interest.

### Summary of Updates

We have generated a GFP-tagged KIN-20 to determine localization of this protein. We showed LIN-42 and KIN-20 co-localize and we added data demonstrating that mutation of the LIN-42 CKBD reduces KIN-20 nuclear localization.

## References

1. Takahashi, J.S. (2017). Transcriptional architecture of the mammalian circadian clock. Nat. Rev. Genet. 18, 164–179.

2. Gallego, M., and Virshup, D.M. (2007). Post-translational modifications regulate the ticking of the circadian clock. Nat. Rev. Mol. Cell Biol. 8, 139–148.

3. Gekakis, N., Staknis, D., Nguyen, H.B., Davis, F.C., Wilsbacher, L.D., King, D.P., Takahashi, J.S., and Weitz, C.J. (1998). Role of the CLOCK protein in the mammalian circadian mechanism. Science 280, 1564–1569.

4. Lee, C., Etchegaray, J.P., Cagampang, F.R., Loudon, A.S., and Reppert, S.M. (2001). Posttranslational mechanisms regulate the mammalian circadian clock. Cell 107, 855–867.

5. Lowrey, P.L., and Takahashi, J.S. (2011). Genetics of circadian rhythms in Mammalian model organisms. Adv. Genet. 74, 175–230.

6. Preußner, M., and Heyd, F. (2016). Post-transcriptional control of the mammalian circadian clock: implications for health and disease. Pflugers Arch. 468, 983–991.

7. Lee, C., Weaver, D.R., and Reppert, S.M. (2004). Direct Association between Mouse PERIOD and CKIɛ Is Critical for a Functioning Circadian Clock. Mol Cell Biol 24, 584–594. 10.1128/MCB.24.2.584-594.2004.

8. Aryal, R.P., Kwak, P.B., Tamayo, A.G., Gebert, M., Chiu, P.-L., Walz, T., and Weitz, C.J. (2017). Macromolecular Assemblies of the Mammalian Circadian Clock. Mol Cell 67, 770–782.e6. 10.1016/j.molcel.2017.07.017.

9. Cao, X., Yang, Y., Selby, C.P., Liu, Z., and Sancar, A. (2021). Molecular mechanism of the repressive phase of the mammalian circadian clock. Proc Natl Acad Sci U S A 118, e2021174118. 10.1073/pnas.2021174118.

10. Akashi, M., Tsuchiya, Y., Yoshino, T., and Nishida, E. (2002). Control of Intracellular Dynamics of Mammalian Period Proteins by Casein Kinase I ɛ (CKIɛ) and CKIδ in Cultured Cells. Mol Cell Biol 22, 1693–1703. 10.1128/MCB.22.6.1693-1703.2002.

11. Eide, E.J., Woolf, M.F., Kang, H., Woolf, P., Hurst, W., Camacho, F., Vielhaber, E.L., Giovanni, A., and Virshup, D.M. (2005). Control of mammalian circadian rhythm by CKIepsilon-regulated proteasome-mediated PER2 degradation. Mol. Cell. Biol. 25, 2795– 2807.

12. Shirogane, T., Jin, J., Ang, X.L., and Harper, J.W. (2005). SCFbeta-TRCP controls clock-dependent transcription via casein kinase 1-dependent degradation of the mammalian period-1 (Per1) protein. J Biol Chem 280, 26863–26872. 10.1074/jbc.M502862200.

13. Narasimamurthy, R., Hunt, S.R., Lu, Y., Fustin, J.-M., Okamura, H., Partch, C.L., Forger, D.B., Kim, J.K., and Virshup, D.M. (2018). CK1δ/ε protein kinase primes the PER2 circadian phosphoswitch. Proc Natl Acad Sci U S A 115, 5986–5991. 10.1073/pnas.1721076115.

14. Philpott, J.M., Freeberg, A.M., Park, J., Lee, K., Ricci, C.G., Hunt, S.R., Narasimamurthy, R., Segal, D.H., Robles, R., Cai, Y., et al. (2023). PERIOD phosphorylation leads to feedback inhibition of CK1 activity to control circadian period. Molecular Cell 83, 1677–1692.e8. 10.1016/j.molcel.2023.04.019.

15. Philpott, J.M., Narasimamurthy, R., Ricci, C.G., Freeberg, A.M., Hunt, S.R., Yee, L.E., Pelofsky, R.S., Tripathi, S., Virshup, D.M., and Partch, C.L. (2020). Casein kinase 1 dynamics underlie substrate selectivity and the PER2 circadian phosphoswitch. eLife 9, e52343. 10.7554/eLife.52343.

16. Xu, Y., Toh, K.L., Jones, C.R., Shin, J.-Y., Fu, Y.-H., and Ptácek, L.J. (2007). Modeling of a human circadian mutation yields insights into clock regulation by PER2. Cell 128, 59–70. 10.1016/j.cell.2006.11.043.

17. Jones, C.R., Campbell, S.S., Zone, S.E., Cooper, F., DeSano, A., Murphy, P.J., Jones, B., Czajkowski, L., and Ptček, L.J. (1999). Familial advanced sleep-phase syndrome: A short-period circadian rhythm variant in humans. Nat Med 5, 1062–1065. 10.1038/12502.

18. Toh, K.L., Jones, C.R., He, Y., Eide, E.J., Hinz, W.A., Virshup, D.M., Ptácek, L.J., and Fu, Y.H. (2001). An hPer2 phosphorylation site mutation in familial advanced sleep phase syndrome. Science 291, 1040–1043. 10.1126/science.1057499.

19. Jeon, M., Gardner, H.F., Miller, E.A., Deshler, J., and Rougvie, A.E. (1999). Similarity of the *C. elegans* Developmental Timing Protein LIN-42 to Circadian Rhythm Proteins. Science 286, 1141–1146. 10.1126/science.286.5442.1141.

20. Lamberti, M.L., Spangler, R.K., Cerdeira, V., Ares, M., Rivollet, L., Ashley, G.E., Coronado, A.R., Tripathi, S., Spiousas, I., Ward, J.D., et al. (2024). Clock gene homologs *lin-42* and *kin-20* regulate circadian rhythms in *C. elegans*. Sci Rep 14, 12936. 10.1038/s41598-024-62303-9.

21. Migliori, M.L., Goya, M.E., Lamberti, M.L., Silva, F., Rota, R., Bénard, C., and Golombek, D.A. (2023). *Caenorhabditis elegans* as a Promising Model Organism in Chronobiology. J Biol Rhythms, 07487304221143483. 10.1177/07487304221143483.

22. Tennessen, J.M., Gardner, H.F., Volk, M.L., and Rougvie, A.E. (2006). Novel heterochronic functions of the *Caenorhabditis elegans* period-related protein LIN-42. Dev Biol 289, 30–43. 10.1016/j.ydbio.2005.09.044.

23. Gissendanner, C.R., Crossgrove, K., Kraus, K.A., Maina, C.V., and Sluder, A.E. (2004). Expression and function of conserved nuclear receptor genes in *Caenorhabditis elegans*. Dev Biol 266, 399–416. 10.1016/j.ydbio.2003.10.014.

24. Abrahante, J.E., Miller, E.A., and Rougvie, A.E. (1998). Identification of Heterochronic Mutants in *Caenorhabditis elegans*: Temporal Misexpression of a Collagen::Green Fluorescent Protein Fusion Gene. Genetics 149, 1335–1351. 10.1093/genetics/149.3.1335.

25. Liu, Z. (1990). Genetic Control of Stage-specific Developmental Events in C. elegans.

26. Banerjee, D., Kwok, A., Lin, S.-Y., and Slack, F.J. (2005). Developmental timing in *C. elegans* is regulated by *kin-20* and *tim-1*, homologs of core circadian clock genes. Dev Cell 8, 287–295. 10.1016/j.devcel.2004.12.006.

27. Hasegawa, K., Saigusa, T., and Tamai, Y. (2005). *Caenorhabditis elegans* opens up new insights into circadian clock mechanisms. Chronobiol. Int. 22, 1–19.

28. Kostrouchova, M., Krause, M., Kostrouch, Z., and Rall, J.E. (1998). CHR3: a *Caenorhabditis elegans* orphan nuclear hormone receptor required for proper epidermal development and molting. Development (Cambridge, England) 125, 1617–1626.

29. Ambros, V. (1989). A hierarchy of regulatory genes controls a larva-to-adult developmental switch in *C. elegans*. Cell 57, 49–57.

30. Ambros, V., and Horvitz, H.R. (1984). Heterochronic mutants of the nematode *Caenorhabditis elegans*. Science 226, 409–416.

31. McCulloch, K.A., and Rougvie, A.E. (2014). *Caenorhabditis elegans period* homolog *lin-42* regulates the timing of heterochronic miRNA expression. Proceedings of the National Academy of Sciences 111, 15450–15455. 10.1073/pnas.1414856111.

32. Perales, R., King, D.M., Aguirre-Chen, C., and Hammell, C.M. (2014). LIN-42, the *Caenorhabditis elegans* PERIOD homolog, Negatively Regulates MicroRNA Transcription. PLoS Genet 10, e1004486. 10.1371/journal.pgen.1004486.s008.

33. Van Wynsberghe, P.M., Finnegan, E.F., Stark, T., Angelus, E.P., Homan, K.E., Yeo, G.W., and Pasquinelli, A.E. (2014). The Period protein homolog LIN-42 negatively regulates microRNA biogenesis in *C. elegans*. Dev Biol 390, 126–135. 10.1016/j.ydbio.2014.03.017.

34. Lažetić, V., and Fay, D.S. (2017). Molting in *C. elegans*. Worm 6, 1–20. 10.1080/21624054.2017.1330246.

35. Monsalve, G.C., Van Buskirk, C., and Frand, A.R. (2011). LIN-42/PERIOD controls cyclical and developmental progression of *C. elegans* molts. Curr Biol 21, 2033–2045. 10.1016/j.cub.2011.10.054.

36. Olmedo, M., Geibel, M., Artal-Sanz, M., and Merrow, M. (2015). A High-Throughput Method for the Analysis of Larval Developmental Phenotypes in *Caenorhabditis elegans*. Genetics 201, 443. 10.1534/genetics.115.179242.

37. Kinney, B., Sahu, S., Stec, N., Hills-Muckey, K., Adams, D.W., Wang, J., Jaremko, M., Joshua-Tor, L., Keil, W., and Hammell, C.M. (2023). A circadian-like gene network programs the timing and dosage of heterochronic miRNA transcription during *C. elegans* development. Dev Cell 58, 2563–2579.e8. 10.1016/j.devcel.2023.08.006.

38. Rhodehouse, K., Cascino, K., Aseltine, L., Padula, A., Weinstein, R., Spina, J.S., Olivero, C.E., and Van Wynsberghe, P.M. (2018). The Doubletime Homolog KIN-20 Mainly Regulates *let-7* Independently of Its Effects on the Period Homolog LIN-42 in *Caenorhabditis elegans*. G3 (Bethesda) 8, 2617–2629. 10.1534/g3.118.200392.

39. Edelman, T.L.B., McCulloch, K.A., Barr, A., Frøkjær-Jensen, C., Jorgensen, E.M., and Rougvie, A.E. (2016). Analysis of a *lin-42/period* Null Allele Implicates All Three Isoforms in Regulation of *Caenorhabditis elegans* Molting and Developmental Timing. G3 (Bethesda) 6, 4077–4086. 10.1534/g3.116.034165.

40. Meeuse, M.W., Hauser, Y.P., Morales Moya, L.J., Hendriks, G., Eglinger, J., Bogaarts, G., Tsiairis, C., and Großhans, H. (2020). Developmental function and state transitions of a gene expression oscillator in *Caenorhabditis elegans*. Mol Syst Biol 16, e9975. 10.15252/msb.20209498.

41. Rüegger, S., Miki, T.S., Hess, D., and Großhans, H. (2015). The ribonucleotidyl transferase USIP-1 acts with SART3 to promote U6 snRNA recycling. Nucleic Acids Res 43, 3344– 3357. 10.1093/nar/gkv196.

42. Hendriks, G.-J., Gaidatzis, D., Aeschimann, F., and Großhans, H. (2014). Extensive oscillatory gene expression during *C. elegans* larval development. Mol Cell 53, 380–392. 10.1016/j.molcel.2013.12.013.

43. Crosby, P., Goularte, N.F., Sharma, D., Chen, E., Parico, G.C.G., Philpott, J.M., Harold, R., Gustafson, C.L., and Partch, C.L. (2023). CHRONO participates in multi-modal repression of circadian transcriptional complexes. Preprint at bioRxiv, 10.1101/2022.10.04.510902 https://doi.org/10.1101/2022.10.04.510902.

44. Dahlberg, C.L., Nguyen, E.Z., Goodlett, D., and Kimelman, D. (2009). Interactions between Casein Kinase Iε (CKIε) and Two Substrates from Disparate Signaling Pathways Reveal Mechanisms for Substrate-Kinase Specificity. PLOS ONE 4, e4766. 10.1371/journal.pone.0004766.

45. LaBella, M.L., Hujber, E.J., Moore, K.A., Rawson, R.L., Merrill, S.A., Allaire, P.D., Ailion, M., Hollien, J., Bastiani, M.J., and Jorgensen, E.M. (2020). Casein Kinase 1δ Stabilizes Mature Axons by Inhibiting Transcription Termination of Ankyrin. Dev Cell 52, 88–103.e6. 10.1016/j.devcel.2019.12.005.

46. An, Y., Yuan, B., Xie, P., Gu, Y., Liu, Z., Wang, T., Li, Z., Xu, Y., and Liu, Y. (2022). Decoupling PER phosphorylation, stability and rhythmic expression from circadian clock function by abolishing PER-CK1 interaction. Nat Commun 13, 3991. 10.1038/s41467-022-31715-4.

47. Lowrey, P.L., Shimomura, K., Antoch, M.P., Yamazaki, S., Zemenides, P.D., Ralph, M.R., Menaker, M., and Takahashi, J.S. (2000). Positional syntenic cloning and functional characterization of the mammalian circadian mutation tau. Science 288, 483–492. 10.1126/science.288.5465.483.

48. Xu, Y., Padiath, Q.S., Shapiro, R.E., Jones, C.R., Wu, S.C., Saigoh, N., Saigoh, K., Ptácek, L.J., and Fu, Y.-H. (2005). Functional consequences of a CKIdelta mutation causing familial advanced sleep phase syndrome. Nature 434, 640–644. 10.1038/nature03453.

49. Marzoll, D., Serrano, F.E., Shostak, A., Schunke, C., Diernfellner, A.C.R., and Brunner, M. (2022). Casein kinase 1 and disordered clock proteins form functionally equivalent, phospho-based circadian modules in fungi and mammals. Proc. Natl. Acad. Sci. U. S. A. 119.

50. Johnson, L.N., Noble, M.E., and Owen, D.J. (1996). Active and inactive protein kinases: structural basis for regulation. Cell 85, 149–158. 10.1016/s0092-8674(00)81092-2.

51. Nolen, B., Taylor, S., and Ghosh, G. (2004). Regulation of Protein Kinases: Controlling Activity through Activation Segment Conformation. Molecular Cell 15, 661–675. 10.1016/j.molcel.2004.08.024.

52. Berger, S., Spiri, S., deMello, A., and Hajnal, A. (2021). Microfluidic-based imaging of complete *Caenorhabditis elegans* larval development. Development 148, dev199674. 10.1242/dev.199674.

53. Zheng, B., Larkin, D.W., Albrecht, U., Sun, Z.S., Sage, M., Eichele, G., Lee, C.C., and Bradley, A. (1999). The *mPer2* gene encodes a functional component of the mammalian circadian clock. Nature 400, 169–173. 10.1038/22118.

54. Kim, E.Y., Ko, H.W., Yu, W., Hardin, P.E., and Edery, I. (2007). A DOUBLETIME kinase binding domain on the *Drosophila* PERIOD protein is essential for its hyperphosphorylation, transcriptional repression, and circadian clock function. Mol Cell Biol 27, 5014–5028. 10.1128/MCB.02339-06.

55. Nawathean, P., Stoleru, D., and Rosbash, M. (2007). A small conserved domain of *Drosophila* PERIOD is important for circadian phosphorylation, nuclear localization, and transcriptional repressor activity. Mol Cell Biol 27, 5002–5013. 10.1128/MCB.02338-06.

56. Cullati, S.N., Chaikuad, A., Chen, J.-S., Gebel, J., Tesmer, L., Zhubi, R., Navarrete-Perea, J., Guillen, R.X., Gygi, S.P., Hummer, G., et al. (2022). Kinase domain autophosphorylation rewires the activity and substrate specificity of CK1 enzymes. Mol Cell 82, 2006–2020.e8. 10.1016/j.molcel.2022.03.005.

57. Gebel, J., Tuppi, M., Chaikuad, A., Hötte, K., Schröder, M., Schulz, L., Löhr, F., Gutfreund, N., Finke, F., Henrich, E., et al. (2020). p63 uses a switch-like mechanism to set the threshold for induction of apoptosis. Nat Chem Biol 16, 1078–1086. 10.1038/s41589-020-0600-3.

58. Harold, R.L., Tulsian, N.K., Narasimamurthy, R., Yaitanes, N., Ayala Hernandez, M.G., Lee, H.-W., Crosby, P., Tripathi, S.M., Virshup, D.M., and Partch, C.L. (2024). Isoform-specific C-terminal phosphorylation drives autoinhibition of Casein kinase 1. Proc Natl Acad Sci U S A 121, e2415567121. 10.1073/pnas.2415567121.

59. Chiou, Y.-Y., Yang, Y., Rashid, N., Ye, R., Selby, C.P., and Sancar, A. (2016). Mammalian Period represses and de-represses transcription by displacing CLOCK-BMAL1 from promoters in a Cryptochrome-dependent manner. Proc Natl Acad Sci U S A 113, E6072– E6079. 10.1073/pnas.1612917113.

60. Narasimamurthy, R., and Virshup, D.M. (2021). The phosphorylation switch that regulates ticking of the circadian clock. Mol Cell 81, 1133–1146. 10.1016/j.molcel.2021.01.006.

61. Brenner, S. (1974). The genetics of *Caenorhabditis elegans*. Genetics 77, 71–94.

62. Church, D.L., Guan, K.L., and Lambie, E.J. (1995). Three genes of the MAP kinase cascade, *mek-2*, *mpk-1/sur-1* and *let-60 ras*, are required for meiotic cell cycle progression in *Caenorhabditis elegans*. Development 121, 2525–2535.

63. Ragle, J.M., Levenson, M.T., Clancy, J.C., Vo, A.A., Pham, V., and Ward, J.D. (2022). The conserved, secreted protease inhibitor MLT-11 is necessary for *C. elegans* molting and embryogenesis. Preprint at bioRxiv, 10.1101/2022.06.29.498124 https://doi.org/10.1101/2022.06.29.498124.

64. Frøkjaer-Jensen, C., Davis, M.W., Hopkins, C.E., Newman, B.J., Thummel, J.M., Olesen, S.-P., Grunnet, M., and Jorgensen, E.M. (2008). Single-copy insertion of transgenes in *Caenorhabditis elegans*. Nat Genet 40, 1375–1383. 10.1038/ng.248.

65. Noma, K., Goncharov, A., Ellisman, M.H., and Jin, Y. (2017). Microtubule-dependent ribosome localization in *C. elegans neurons*. Elife 6, e26376. 10.7554/eLife.26376.

66. Frøkjær-Jensen, C., Davis, M.W., Ailion, M., and Jorgensen, E.M. (2012). Improved Mos1-mediated transgenesis in *C. elegans*. Nat Meth 9, 117–118. 10.1038/nmeth.1865.

67. Gudipati, R.K., Braun, K., Gypas, F., Hess, D., Schreier, J., Carl, S.H., Ketting, R.F., and Großhans, H. (2021). Protease-mediated processing of Argonaute proteins controls small RNA association. Mol Cell 81, 2388–2402.e8. 10.1016/j.molcel.2021.03.029.

68. Schindelin, J., Arganda-Carreras, I., Frise, E., Kaynig, V., Longair, M., Pietzsch, T., Preibisch, S., Rueden, C., Saalfeld, S., Schmid, B., et al. (2012). Fiji: an open-source platform for biological-image analysis. Nat Methods 9, 676–682. 10.1038/nmeth.2019.

69. Chiu, C.-L., Clack, N., and Community, T.N. (2022). napari: a Python Multi-Dimensional Image Viewer Platform for the Research Community. Microscopy and Microanalysis 28, 1576–1577. 10.1017/S1431927622006328.

70. Sheffield, P., Garrard, S., and Derewenda, Z. (1999). Overcoming expression and purification problems of RhoGDI using a family of “parallel” expression vectors. Protein Expr Purif 15, 34–39. 10.1006/prep.1998.1003.

71. Fribourgh, J.L., Srivastava, A., Sandate, C.R., Michael, A.K., Hsu, P.L., Rakers, C., Nguyen, L.T., Torgrimson, M.R., Parico, G.C.G., Tripathi, S., et al. (2020). Dynamics at the serine loop underlie differential affinity of cryptochromes for CLOCK:BMAL1 to control circadian timing. Elife 9, e55275. 10.7554/eLife.55275.

72. Parico, G.C.G., Perez, I., Fribourgh, J.L., Hernandez, B.N., Lee, H.-W., and Partch, C.L. (2020). The human CRY1 tail controls circadian timing by regulating its association with CLOCK:BMAL1. Proc Natl Acad Sci U S A 117, 27971–27979. 10.1073/pnas.1920653117.

73. Frankenfield, A.M., Ni, J., Ahmed, M., and Hao, L. (2022). Protein Contaminants Matter: Building Universal Protein Contaminant Libraries for DDA and DIA Proteomics. J Proteome Res 21, 2104–2113. 10.1021/acs.jproteome.2c00145.

