## Supplementary material for "A conserved chronobiological complex times *C. elegans* development": Combined Supplemental Figures

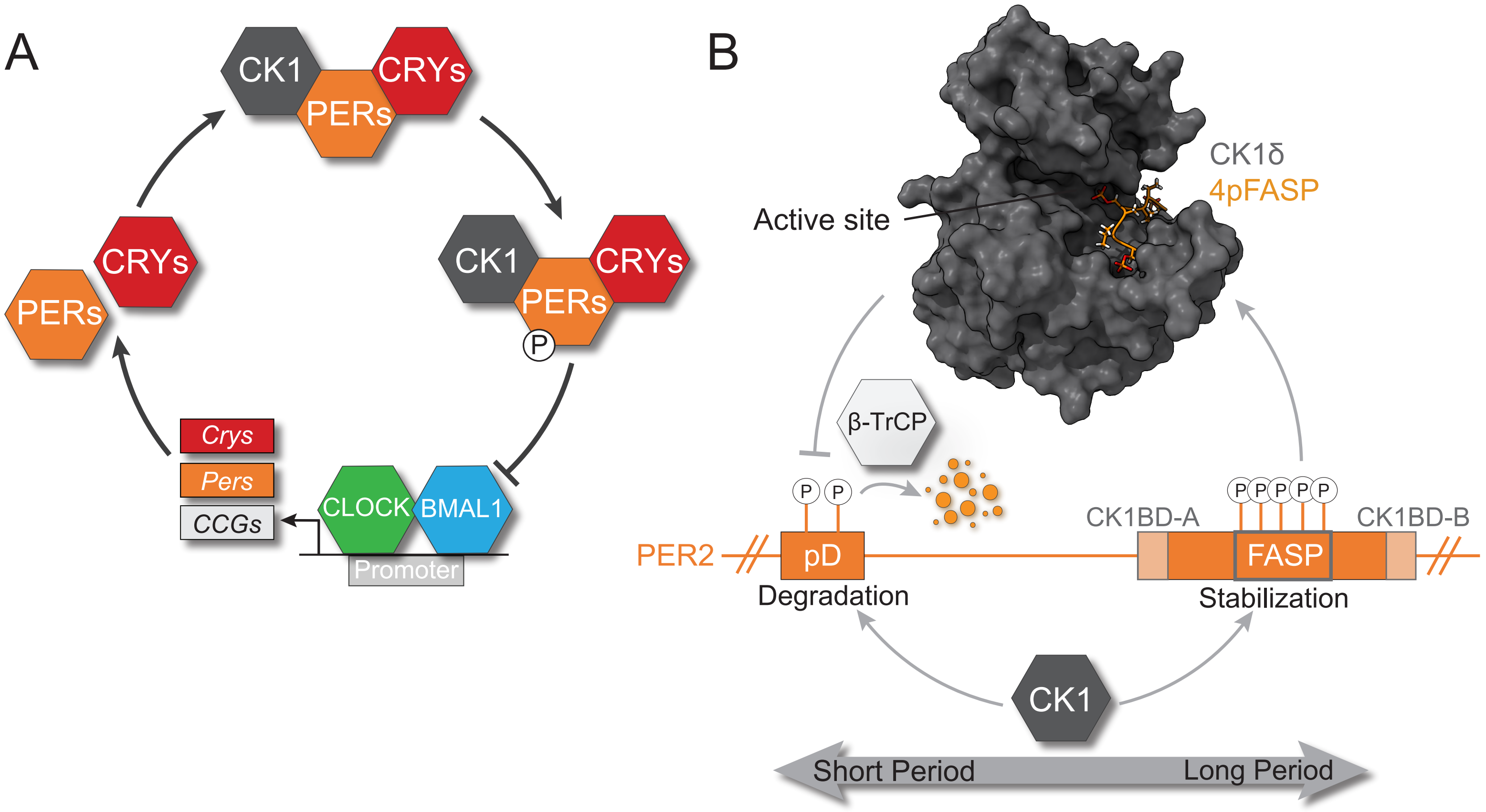


**Figure S1 PER proteins have a crucial role in mammalian circadian rhythms. (A)** Cartoon schematic of the primary transcription-translation feedback loop which generates ~24-hour rhythms in mammals. CCGs, Clock-controlled genes. **(B)** Cartoon schematic illustrating the mammalian phosphoswitch that dictates CK1-dependent regulation of PER stability. Crystal structure of human CK1δ (gray) bound to phosphorylated PER2 FASP (orange, 4pFASP) peptide, PDB: 8d7o. pD, phosphodegron; FASP, Familial Advanced Sleep Phase.

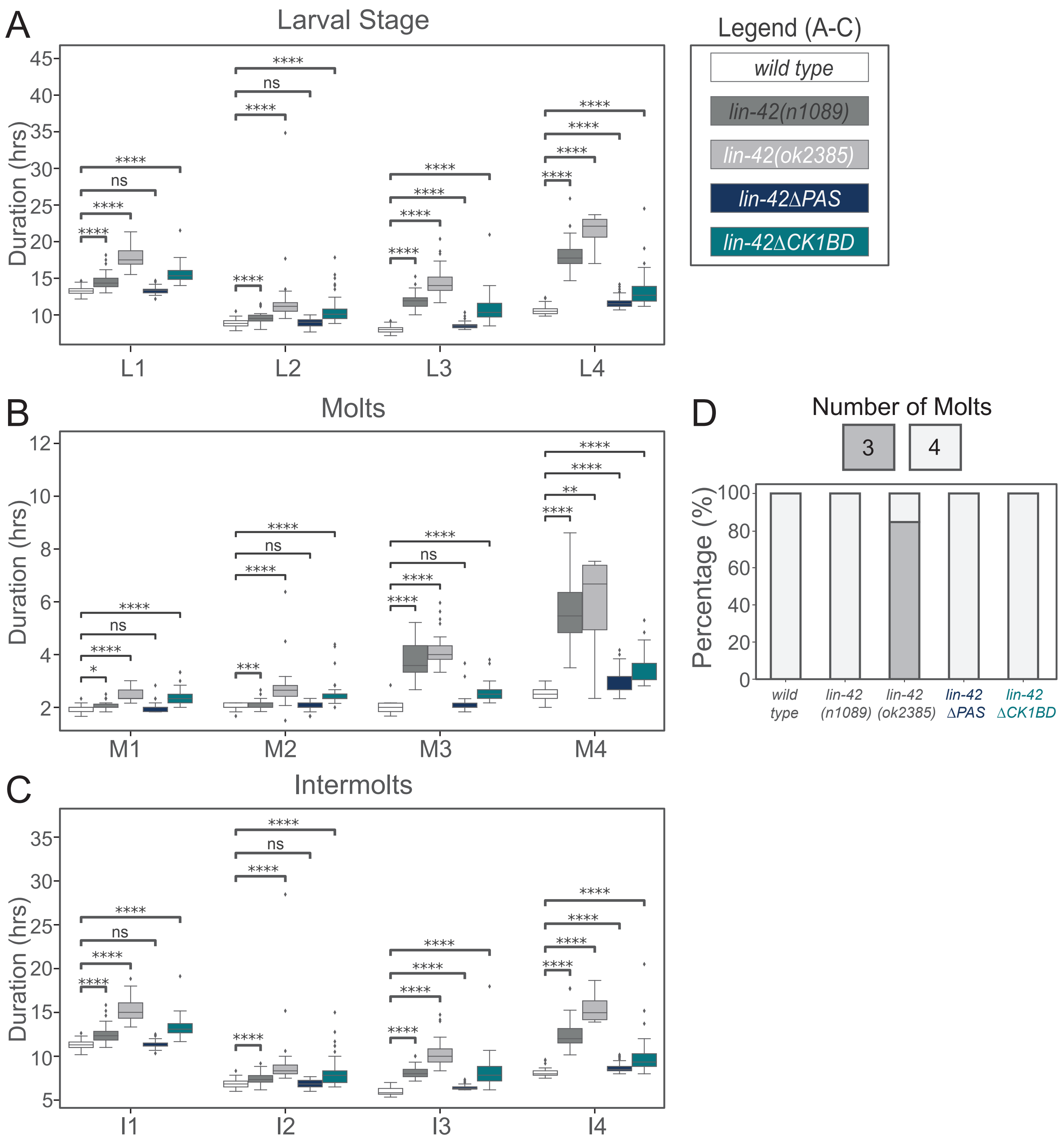


**Figure S2 Larval stage durations for *lin-42* mutant animals.** Boxplots showing durations (in hours) for **(A)** larval stages **(B)** molts and **(C)** intermolts from luciferase assay. Wild type in white, *lin-42(n1089)* in dark grey, *lin-42(ok2385)* in light grey, *lin-42ΔPAS* in blue and *lin-42ΔCK1BD* in green. **(D)** Bar plot showing the number of molts from the luciferase assay of the indicated genotypes.

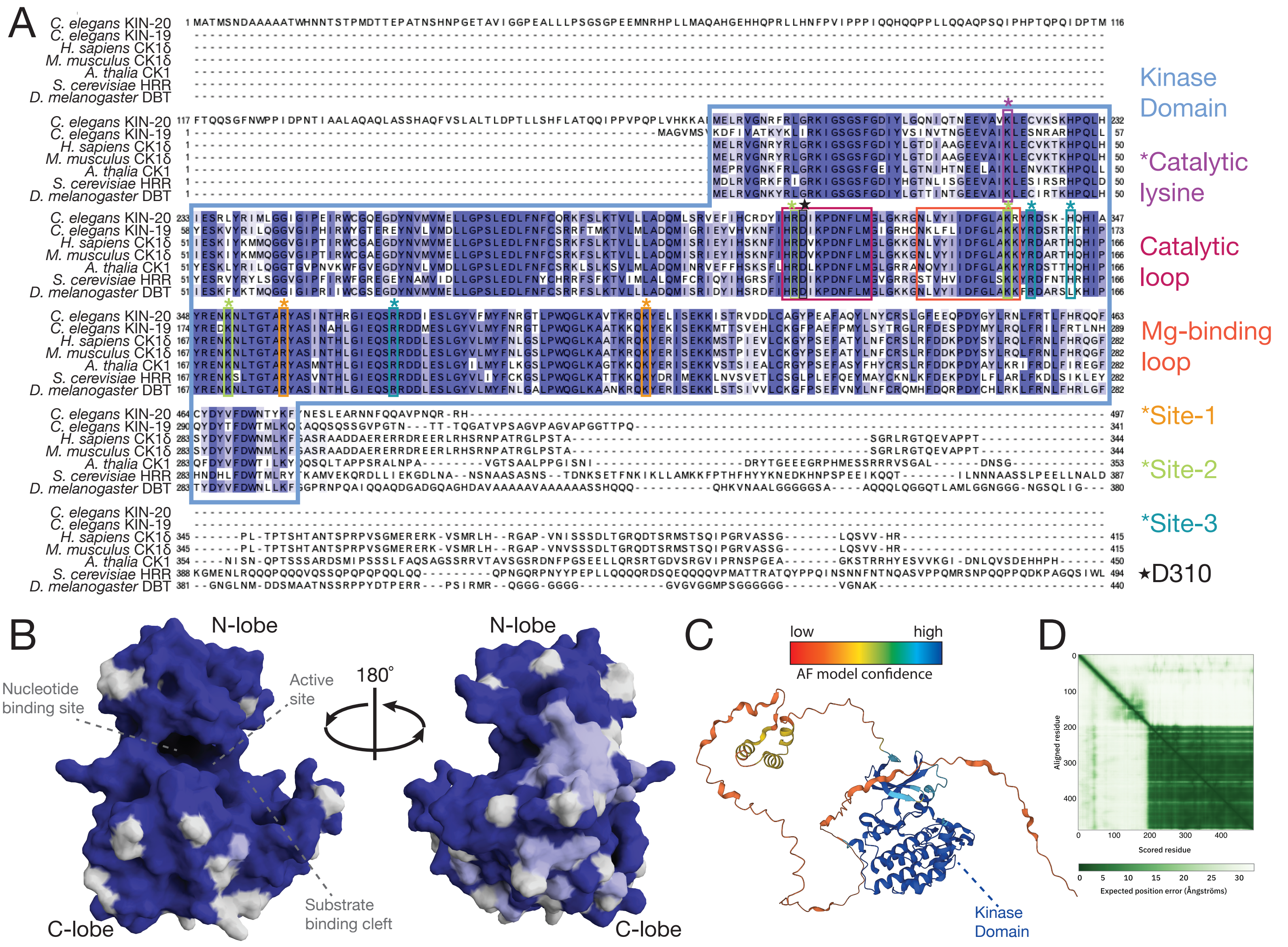


**Figure S3 Conservation and structural prediction of *C. elegans* KIN-20. (A)** Sequence alignment of *C. elegans* KIN-20 and KIN-19, *H. sapiens* CK1δ, *M. musculus* CK1δ, *A. thalia* CK1, *S. cerevisiae* HRR25 (HRR), and *D. melanogaster* Doubletime (DBT). Important enzymatic sequence features are boxed; full kinase domain (blue box - 79% identical between KIN-20 and human CK1δ), catalytic lysine (purple - conserved in KIN-20), catalytic loop (pink - 100% conserved between KIN-20 and human CK1δ), Magnesium (Mg)-binding loop (orange - 12 out of 13 residues conserved between KIN-20 and human CK1), anion coordination sites 1 (yellow), 2 (green), and 3 (blue) (all conserved in KIN-20). **(B)** Crystal structure of *H. sapiens* CK1δ (PDB 6pxo). Dark blue indicates residue is conserved in KIN-20; residues that diverge are highlighted in pale blue (similar amino acid) or not conserved (white). **(C)** AlphaFold structural prediction of KIN-20 (https://alphafold.ebi.ac.uk/entry/A8X4B3) colored by the model confidence. **(D)** KIN-20 AlphaFold plot of predicted aligned error.

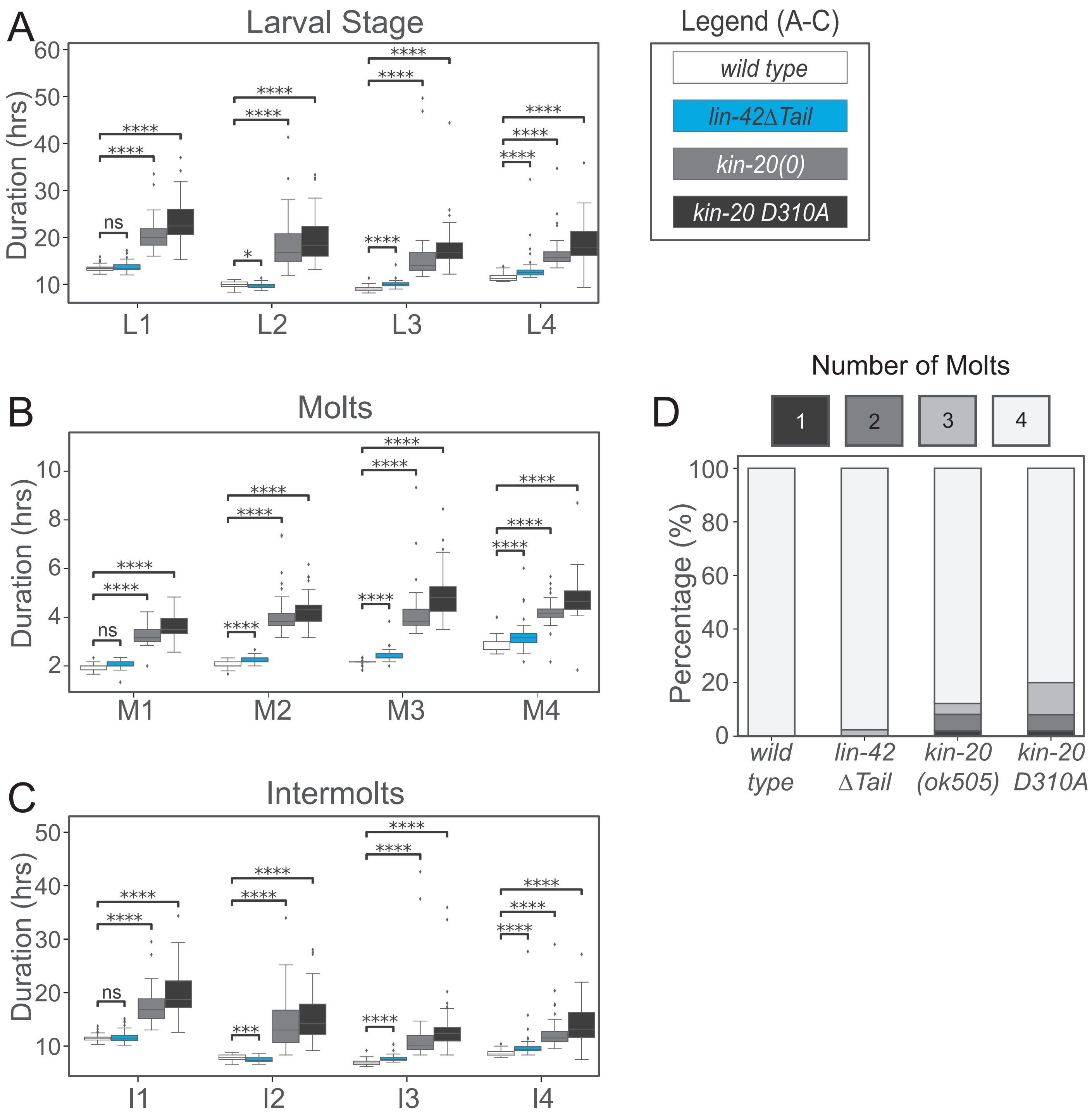


**Figure S4 Larval stage durations for *kin-20* and *lin-42* mutant animals.** Boxplots showing durations (in hours) for **(A)** Larval stages **(B)** Molts **(C)** Intermolts from luciferase assay. Wild type in white, *lin-42(ΔTail)* in blue, *kin-20(0)* in light grey, *kin-20 D310A* in dark grey. Statistics were done using the Mann-Whitney U-test. Stars indicate the significance of difference between the Wt strain and the different lin-42 and kin-20 mutant animals: * $p<0.05$, ** $p<0.01$, *** $p<0.001$, **** $p<0.0001$. **(D)** Bar plot showing the number of molts detected in the assay in percentage of animals.

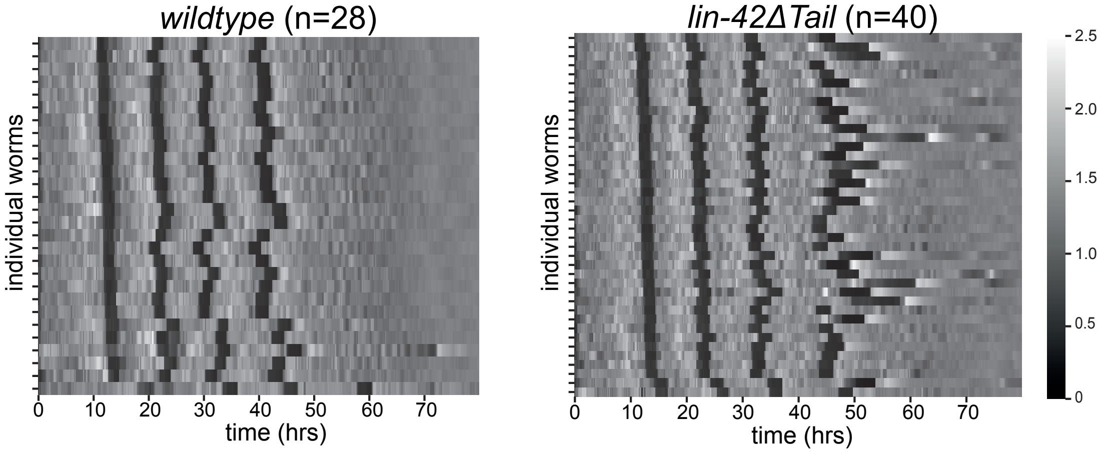


**Figure S5 Luciferase replicate of *lin-42(∆Tail)*.** Heatmaps showing trend-corrected luminescence traces from the indicated genotype. Each horizontal line represents one animal. Traces are sorted to the entry of the first molt. Darker color indicates low luminescence signal and corresponds to the molts.

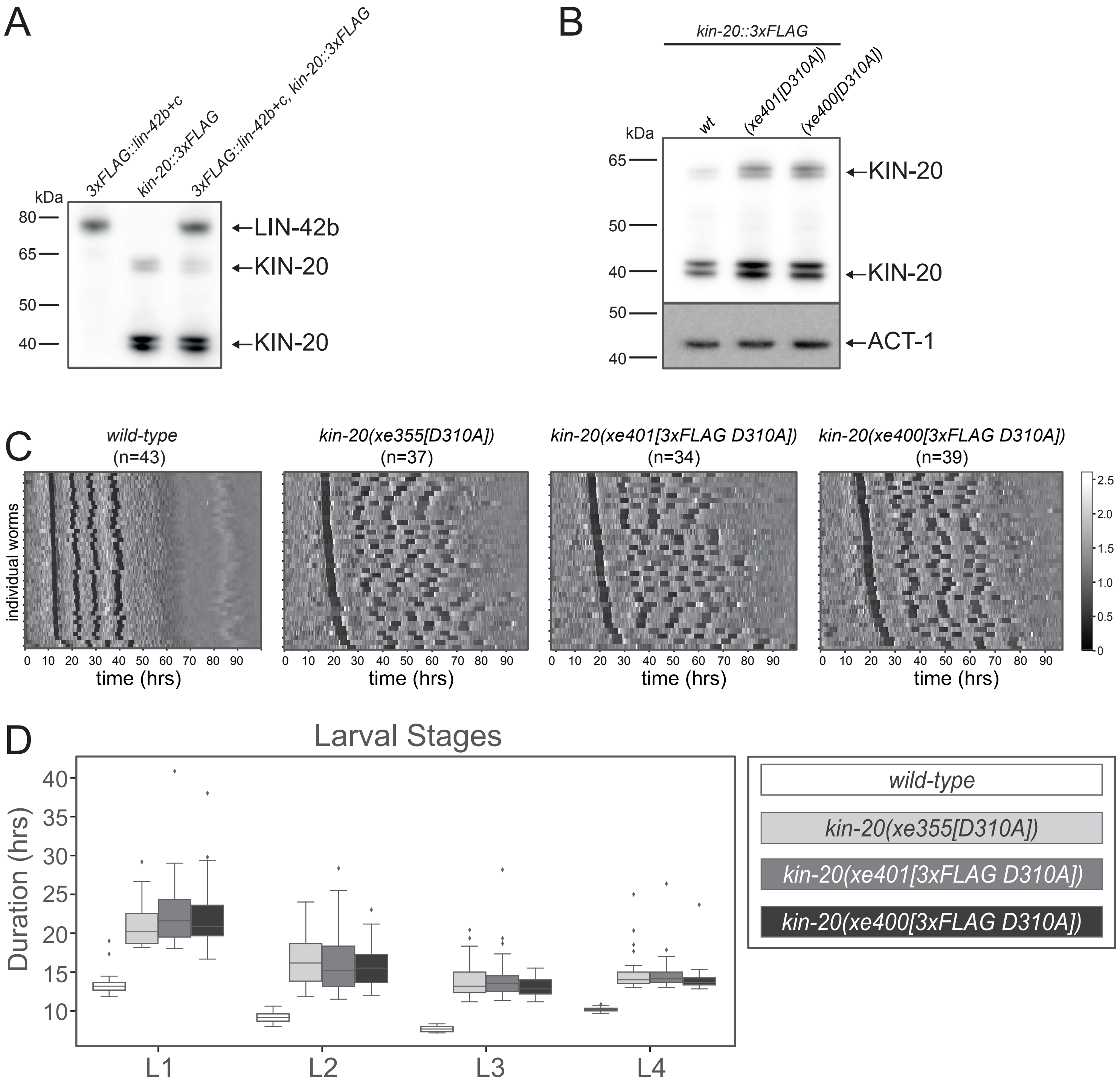


**Figure S6 *kin-20(D310A)* affects molt timing. (A)** Western Blot with extracts from *3xflag::lin-42b+c, 3xflag::kin-20* and *3xflag::lin-42b+c; 3xflag::kin-20* larvae (mid L4 stage). Blot probed with anti-FLAG-HRP (1:1000). Arrows indicate bands for LIN-42b and KIN-20. **(B)** Western Blot with extracts from *3xflag::kin-20* wild-type, *(xe401[D310A])* and *(xe400[D310A])* mutant animals. Top panel probed with anti-FLAG-HRP (1:1000). Lower panel probed with anti-actin-1 (1:7500). Arrows indicate bands for KIN-20. **(C)** Heatmaps showing trend-corrected luminescence traces from the indicated genotype. Each horizontal line represents one animal. Traces are sorted by entry into the first molt. Darker color indicates low luminescence signal and corresponds to the molts. **(D)** Boxplots showing the duration (in hours) for the larval stage from the luciferase assay.

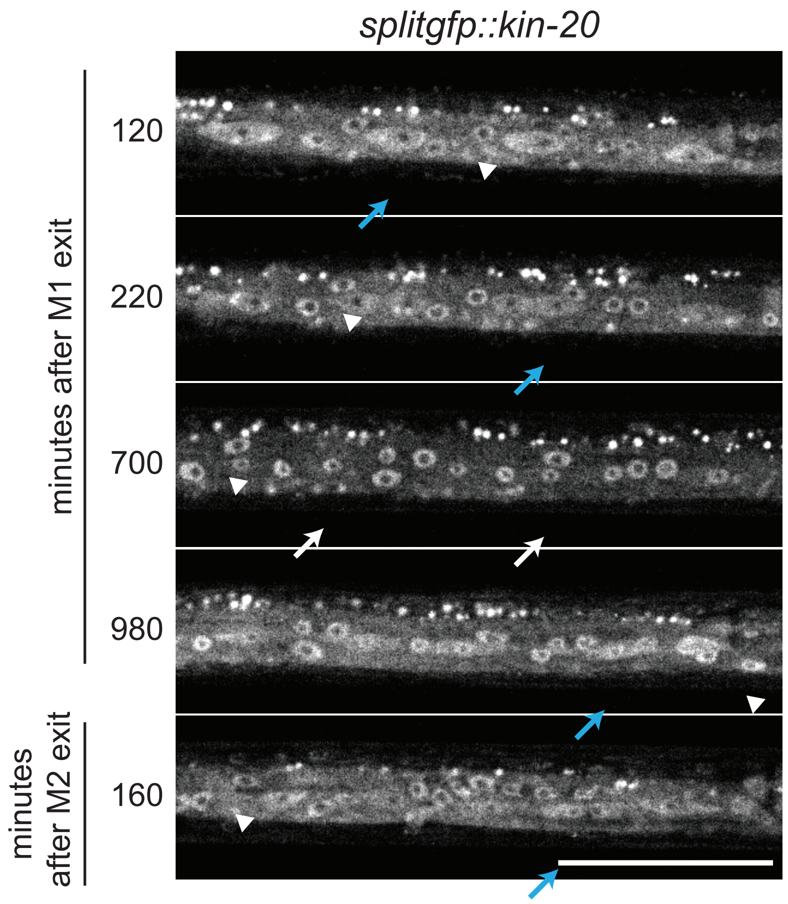


**Figure S7 KIN-20 dynamics during L2 - L3 stage.** Microscopy images of one *splitgfp::kin-20* larva from a microfluidics experiment at indicated timepoints. Time indicated in minutes after Molt 1 (M1) or Molt 2 (M2) exit. Arrows indicate seam cell nuclear (white) and cytoplasmic (blue) localization; white arrowheads indicate nuclear hyp7 localization. Scale bar=50 µm.

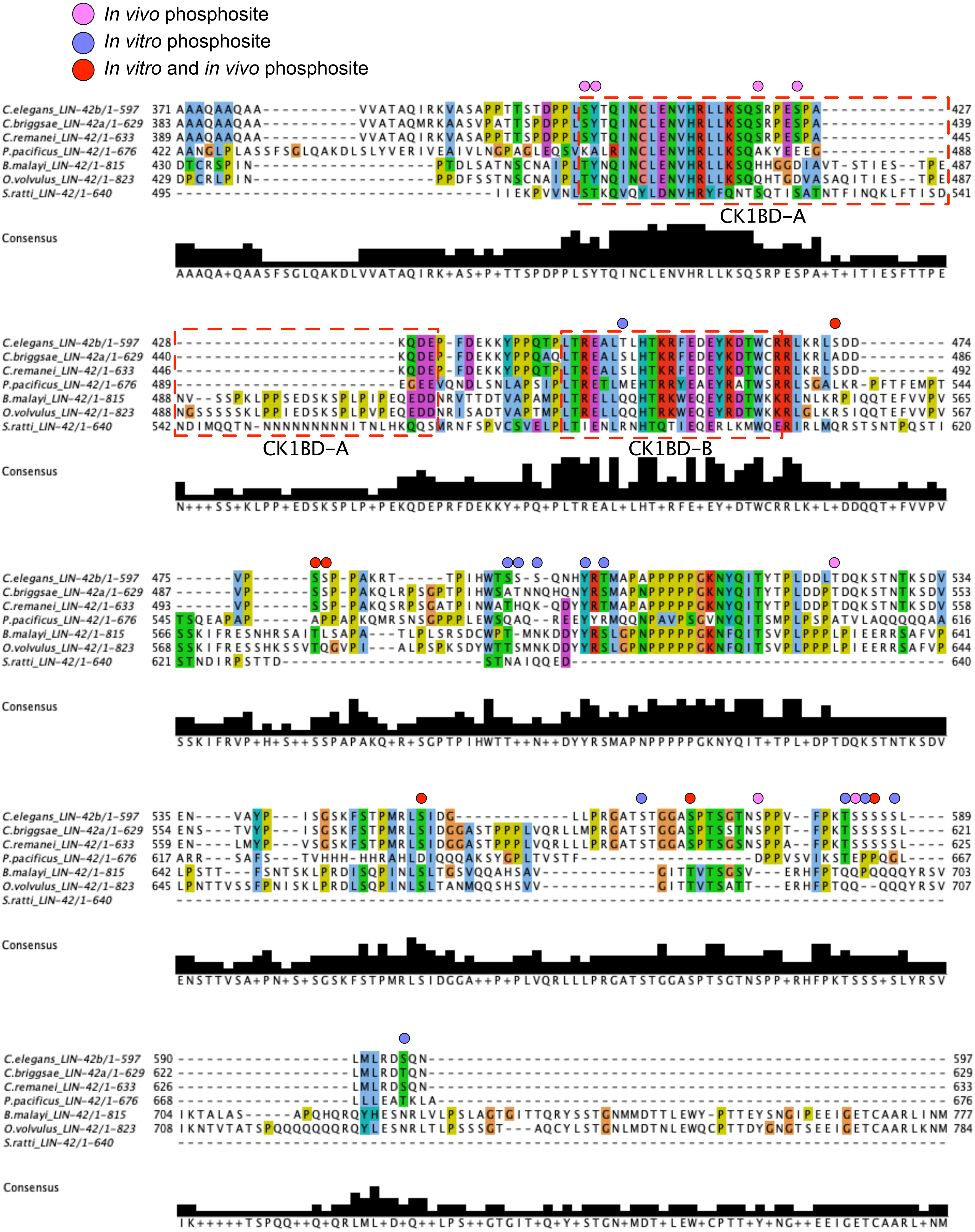


**Figure S8 Alignment of nematode LIN-42 protein sequences.** LIN-42 homologs from the indicated nematode species were aligned using Clustal Omega. The length in amino acids of each homolog follows the species and homolog name. To the left and right of the alignment are amino acid positions of the end residues for each protein. Blue shading indicates conserved sequences and the histogram at the bottom depicts the degree of conservation with a consensus sequence listed below. The positions of the *C. elegans* CK1BD-A and CK1BD-B motifs are indicated. The location of the phosphosites found in our *in vivo*, *in vitro* and both datasets are indicated.
