## Supplemental tables for "A conserved chronobiological complex times *C. elegans* development": Table S1 lin-42 phenotypes.pdf

Table S1 *lin-42* mutant phenotypes

| Strain | Allele | Short name | Arrest[a] (%) | Precocious alae[b] (%) | | | Animals that reached adulthood | |
|---|---|---|---|---|---|---|---|---|
| | | | | Total | Complete | In-complete | BOW[c] (%) | Brood size[d] |
| *N2* | *wild-type* | | 0 | 0 | 0 | 0 | 0 | 313 ± 25 |
| *MT2257* | *lin-42(n1089)* | | 0 | 94.4 | 72.2 | 22.2 | 34.4 | 180 ± 58 |
| *RB1843* | *lin-42(ok2385)* | | 48.1 | 80 | 40 | 40 | 85 | nd* |
| *VC398* | kin-20(ok505) | | 42.5 | 0 | 0 | 0 | 52.2 | 52 ± 21 |
| *JDW580* | *lin-42(wrd182)* | CK1BD-ΔA | 1 | 0 | 0 | 0 | 0 | 318 ± 23 |
| *JDW581* | *lin-42(wrd183)* | CK1BD-ΔA | 1.1 | 0 | 0 | 0 | 0 | 294 ± 32 |
| *JDW664* | *lin-42(wrd238)* | CK1BD -ΔB | 1.5 | 0 | 0 | 0 | 0 | 287 ± 39 |
| *JDW686* | *lin-42(wrd239)* | CK1BD -ΔB | 4.5 | 0 | 0 | 0 | 1.6 | 277 ± 23.4 |
| *JDW335* | *lin-42(wrd63)* | ΔCK1BD | 28.9 | 6.8 | 0 | 6.8 | 1 | 149 ± 41 |
| *JDW577* | *lin-42(wrd179)* | ΔCK1BD | 27.3 | 7.7 | 0 | 7.7 | 5 | 159 ± 31 |
| *JDW439* | *lin-42(wrd107)* | ΔTail | 0 | 56.3 | 5.4 | 50.9 | 29.1 | 261 ± 58 |
| *JDW579* | *lin-42(wrd181)* | ΔTail | 0 | 43.4 | 0 | 43.4 | 18.5 | 270 ± 39 |
| *JDW723* | *lin-42(wrd262)* | ΔCK1BD + Tail | 7.6 | 85.7 | 26.2 | 59.5 | 98.6 | nd* |
| *JDW619* | *lin-42(wrd217)* | ΔCK1BD + Tail | 0 | 85.4 | 43.8 | 41.7 | 97.4 | nd* |
| *JDW340* | *lin-42(wrd67)* | ΔPASA/B | 0 | 13.5 | 0 | 13.5 | 1.5 | 251 ± 38 |
| *JDW648* | *lin-42(wrd227)* | ΔPASA/B | 0 | 24.6 | 1.8 | 22.8 | 0 | 283 ± 30 |

All animals grew from eggs hatched onto seeded plates at low density after sodium hypochlorite treatment. $n \geq 20$ for all analyses. [a]Percentage of animals that failed to reach adulthood by day 8 after hatching or died as young larvae [b]Percentage of animals with alae formation in early L4 (L4.0-L4.2). [c]Percentage of animals that exhibited bag-of-worms phenotype. [d]Average number of progeny from fertile adults; arrested and bagged animals were not included in the calculation.*Broods were not counted in animals with high instance of BOW phenotype
